# Non-canonical Hedgehog signaling through L-type voltage gated Ca^2+^ channels controls CD8^+^ T cell killing

**DOI:** 10.1101/2021.03.01.433424

**Authors:** Joachim Hanna, Chrysa Kapeni, Louise O’Brien, Valentina Carbonaro, Flavio Beke, Chandra Chilamakuri, Maike de la Roche

**Author notes:** Corresponding author. (M.d.l.R.).

## Abstract

Cytotoxic CD8^+^ T lymphocytes (CTLs) are critical to the immune response against intracellular pathogens and cancer and act by eliminating infected and malignant cells through targeted secretion of cytotoxic granules. Hedgehog (Hh) signaling has been shown to be critical for CTL killing. Interestingly, Hh signaling in CD8^+^ T cells is not induced by extracellular Hh ligands but is initiated upon T cell receptor (TCR) engagement. How the TCR induces the Hh pathway independently of extracellular Hh ligands is unknown. Here we show that the Hh transcription factor Gli1 is essential for efficient CTL function and is induced downstream of the TCR by an extracellular Ca^2+^ influx selectively controlled by L-type voltage gated Ca^2+^ channels localized at the plasma membrane. We demonstrate that this novel mode of Hh signaling induction is independent of the canonical Hh pathway and represents the primary mechanism of Gli1 induction in naïve CD8^+^ T cells, while CTLs can also activate Gli1 via MAP Kinase signaling. Importantly, we show that this L-type voltage gated Ca^2+^ channel-controlled Gli1 induction is functionally required for CTL killing in mice and humans. Gli inhibitors are currently in clinical trials against various cancers and our observations indicate that they likely inhibit the anti-tumor response.

**Significance statement:** Cytotoxic CD8^+^ T cells (CTLs) kill infected and malignant cells by targeted secretion of cytotoxic granules. Hedgehog signaling is critical for effective CTL killing and is activated by the T cell receptor (TCR) independently of exogenous Hedgehog ligands. This study shows that Hedgehog transcription factor Gli1 is required for CTL killing and identifies L-type voltage gated Ca^2+^ channels (Cav1) as essential regulators of CTL killing in mouse and human, by virtue of their ability to activate Gli1 downstream of the TCR. This Cav1-Gli1 axis operates independently of canonical Hedgehog signaling. Our work suggests that caution is required when using Gli inhibitors, currently in trials as anti-cancer therapeutics, since they may dampen the anti-tumor response.

## INTRODUCTION

Cytotoxic CD8^+^ T lymphocytes (CTLs) eradicate infected and cancerous cells by targeted release of cytotoxic granules. Two types of immune synapses are important for this to occur: the signaling and the cytotoxic synapse. The signaling synapse initiates CTL differentiation and is formed between a naïve CD8^+^ T cell and an antigen presenting cell (APC). The cytotoxic synapse is formed between a CTL and its target cell whereby engagement of the TCR initiates the secretion of granules containing cytotoxic perforin and granzyme. Both synapses promote optimal TCR signaling ^1^.

Immune synapses are structurally and functionally very similar to the primary cilium, a hairlike protrusion from the cell body present on most cells. For example, both structures dock the centrosome at the plasma membrane via distal appendage proteins and are key signaling hubs and sites of focussed endo- and exocytosis ^1, 2^. The similarities between immune synapses and primary cilia prompted us to study Hedgehog (Hh) signaling – a pathway functionally tied to the primary cilium in vertebrates – at the T cell synapse.

Canonical Hh signaling is initiated when one of three Hh ligands - Sonic Hh (Shh), Indian Hh (Ihh) or Desert Hh (Dhh) - bind to the transmembrane receptor Ptch at the base of the primary cilium. Upon ligand binding, Ptch releases its inhibition of the key signal transducer Smo that translocates to the cilium and activates glioma associated oncogene (Gli) transcription factors (Gli1, Gli2, and Gli3). Gli transcription factors translocate into the nucleus and initiate Hh target gene transcription ^3^. More recently, however, various non-canonical Hh signaling modes have been described which can be divided into two major groups. The first group is independent of Gli transcription: Ptch can act as a dependence receptor and triggers apoptosis in the absence of Hh ligands ^4^, and Smo can regulate the actin cytoskeleton via small GTPases RhoA and Rac1 which occurs through G-proteins and PI3K in fibroblasts and through Src and Fyn in neurons ^5, 6^. Smo can also trigger calcium (Ca^2+^) release from the endoplasmic reticulum (ER) and in spinal neurons through Gi protein and PLC*γ*-catalysed generation of IP3 and the opening of IP3-dependent channels ^7^. The second group is Gli transcription-dependent and Ptch/Smo-independent. This non-canonical activation of Gli1 has been described in cancers and stem cells ^8^. Positive regulators include the MAP Kinase (MAPK) pathway, PI3K-AKT-mTOR, and TGFβ signaling as well as oncogenes such as c-myc.

Although various roles for Hh signaling during T cell development in the thymus have been proposed ^9^, little is known about Hh signaling in mature T cells. We have previously shown that Hh signaling is necessary for CD8^+^ T cell killing and, interestingly, is initiated independently of extracellular Hh ligands ^10^. We previously demonstrated that proximal TCR signaling is the main inducer of the Hh pathway ^10^ but the mechanism by which Hh signaling is initiated downstream of the TCR remains unknown.

Gli1 is the only Gli transcription factor expressed in CD8^+^ T cells. Here we show that Gli1 is critical for CD8^+^ T cell killing. We demonstrate a key role for MAPK signaling inducing Gli1 in CTLs, as has been previously described in other cell types and tumor cells ^11^. Additionally, we find binding sites of the MAPK induced transcription factor activator protein 1 (AP-1) in the Gli1 promoter. Most importantly, we identify a previously unknown, non-canonical mode of Hh signaling that culminates in Gli activation via L-type voltage gated Ca^2+^ channels (Cav1 family) in both CTLs and naïve CD8^+^ T cells.

Published work has shown that constitutive loss of Cav1 channels in all tissues leads to defects in T cell development and maturation, with subsequent functional impairment of peripheral T cells ^12, 13^. However, it is not fully clear what cell-intrinsic role these channels play in fully mature T cells. Our work demonstrates that Cav1 channels are critical for mature CD8^+^ T cell killing by virtue of their ability to induce Gli1 via non-canonical Hh signaling.

## RESULTS

### Gli1 is important for cytotoxic T cell (CTL) killing

Gli1 is the only Gli transcription factor reported to be present in CD8^+^ T cells ^10^ and functions as a reliable marker of Hh signaling activation ^14^. Previous work has shown that CTLs treated with the small molecule Gli inhibitor GANT61 have diminished killing ability ^10^. We wanted to confirm this observation genetically and generated *Gli1^eGFP/eGFP^* (*Gli1* KO) by breeding *Gli1^eGFP/+^* mice to homozygosity, disrupting the expression of the *Gli1* gene. *Gli1* KO mice have a phenotypically normal peripheral T cell compartment (Suppl. Fig. 1) and CD8^+^ T cells from the *Gli1* KO mice lack Gli1 protein (Fig. 1A). CD8^+^ T cells from OTI TCR transgenic mice recognize ovalbumin (Ova) peptide residues 257-264 in the context of MHCI H2K^b^ and we bred *Gli1* KO mice expressing the OTI TCR. We compared *Gli1* WT and KO OTI cells in their ability to kill Ova-presenting EL-4 murine lymphoma target cells. Strikingly, specific T cell killing was reduced by 25-50% in the *Gli1* KO cells compared with the *Gli1* WT controls (Fig. 1B). Thus, Gli1 is functionally important for effective CTL killing.

**Figure 1:**
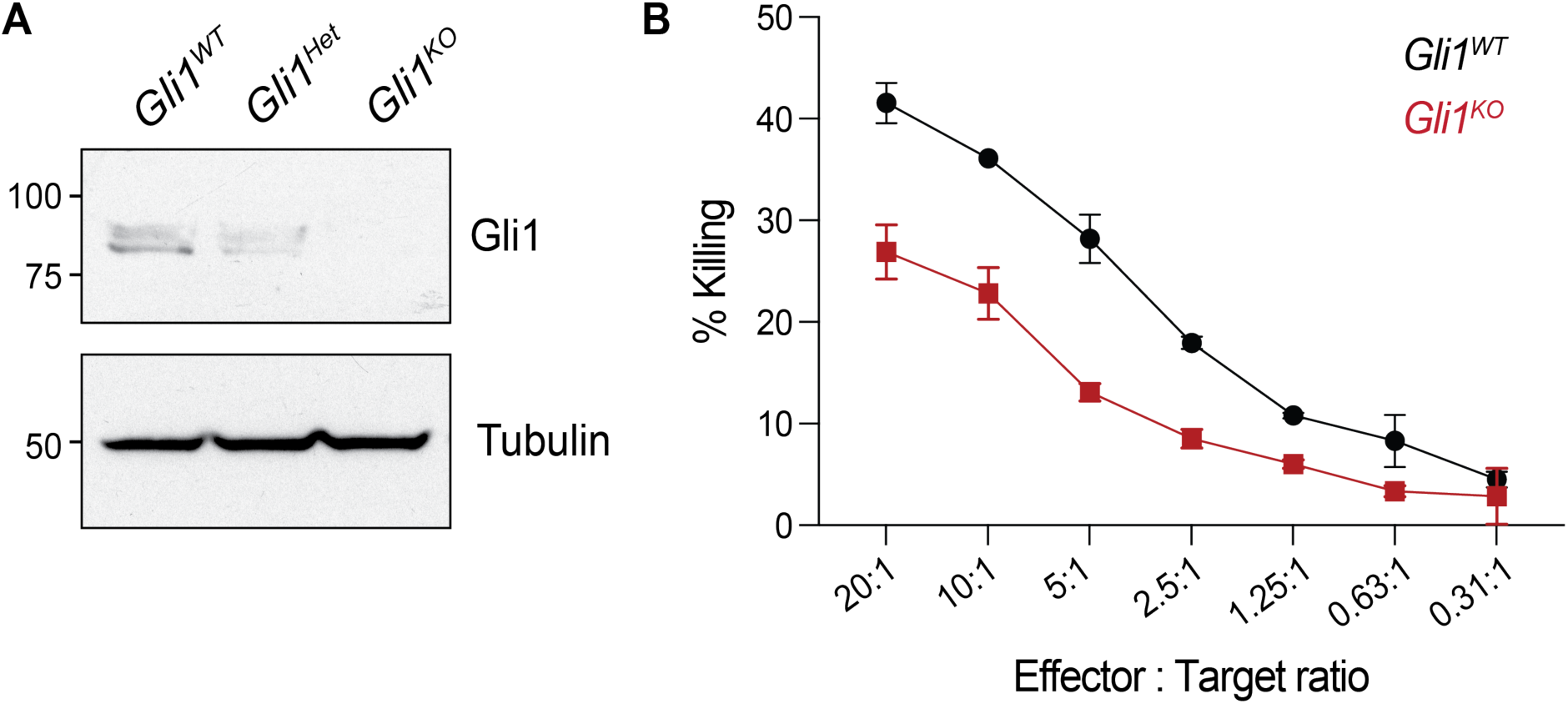
Loss of *Gli1* diminishes CTL killing. (**A**) CTLs were generated from *Gli1^WT^*, *Gli1^Het^* and *Gli1^KO^* mice and restimulated on day 10 with plate-bound anti-CD3 antibodies for 24h. Cells were lysed and lysates blotted for protein expression of Gli1 and tubulin. n=2 independent experiments. Molecular masses are shown in kilodaltons. (**B**) On day 7 post stimulation CTLs from *Gli1^WT^* and *Gli1^KO^* mice were co-cultured with ovalbumin-pulsed EL-4 target cells for 4 hours at the indicated effector to target ratios and subjected to an LDH cytotoxicity assay. Representative data of n=3 independent experiments. Error bars indicate SD.

### MAP kinase signaling downstream of the TCR promotes Gli1 induction in CTLs

CD8^+^ T cell killing is initiated by engagement of the TCR and the TCR-associated proximal tyrosine kinase Lck has been shown to be required for the induction of Gli1 ^10^. Induction of *Gli1* mRNA is a robust readout of Gli1 activation and thus active Hh signalling ^14^. We therefore wanted to investigate how signaling downstream of the TCR leads to Gli1 activation.

TCR engagement by cognate antigen presented on MHCI molecules leads to recruitment of Lck to the TCR complex (Fig. 2A). Lck phosphorylates ZAP70 (ζ-chain-associated protein kinase of 70 kDa) which in turn phosphorylates the adaptor protein LAT (Linker for activation of T cells) leading to the formation of the LAT signalosome. The LAT signalosome propagates signal branching into three major signaling pathways ultimately culminating in the activation of the transcription factors NFAT, NF*κ*B, and AP-1 ^15^.

**Figure 2:**
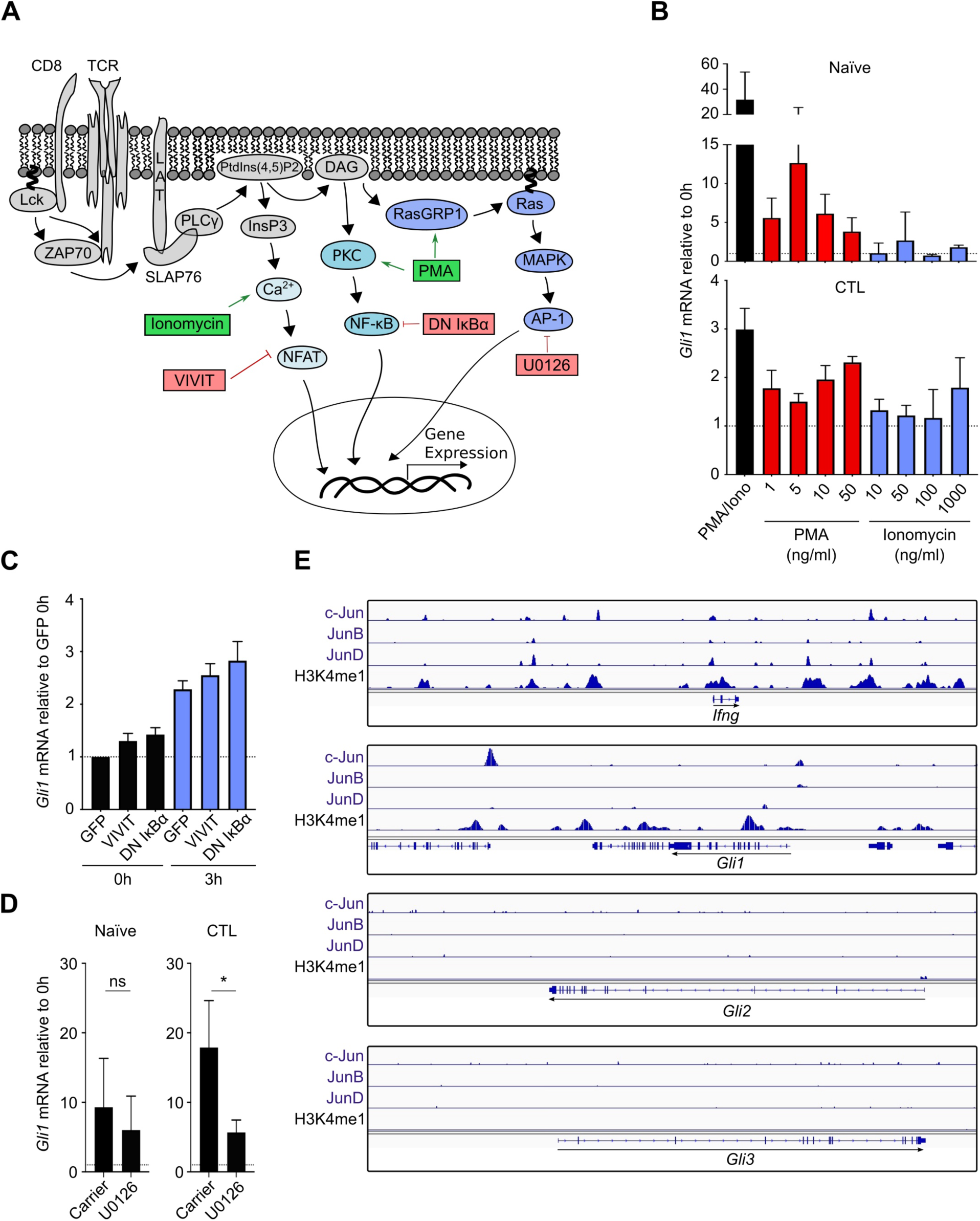
MAPK signalling drives Gli1 induction in CTLs post TCR stimulation. (**A**) Schematic overview of signalling downstream of the TCR culminating in the activation of transcription factors NFAT, NFkB and AP-1. Small molecule agonists (green) as well as antagonists and dominant negative constructs (red) of the branches of TCR signalling are shown. (**B-D**) CD8^+^ T cells were isolated from spleens and inguinal lymph nodes of Rag2^-/-^ OT-I mice. (**B**) Naïve CD8^+^ T cells or CTLs were treated with indicated doses of PMA and/or Ionomycin for 3 hours (naïve, top panel) (CTL, bottom panel) before being subjected to qRT-PCR analysis. For double treatment with PMA and Ionomycin, 50ng/ml PMA and 1µg/ml Ionomycin was used. n=3 independent experiments. (**C**) CTLs were nucleofected with GFP, GFP-VIVIT or Dominant Negative IkBα (DN IkBα) on day 6. Cells were restimulated with anti-CD3 on day 7 for 3 hours for qRT-PCR analysis. n=3 independent experiments. (**D**) Naïve CD8^+^ T cells were stimulated for 6h with plate-bound anti-CD3/CD28 antibodies in the presence of 10µM U0126 or carrier control (left). CTLs were restimulated for 15h with plate-bound anti-CD3 in the presence of 10µM U0126 or carrier control (right) before RNA was extracted for qRT-PCR analysis. n=3 independent experiments. (B-D) Data is normalized to *CD3ε* as a reference gene. Similar results were obtained when *Tbp* was used as a reference gene. Error bars indicate SD. p values were calculated using an unpaired two-tailed Student’s t test. * indicates p<0.05. (**E**) Analysis of ChIP-Seq data from (*Kurachi et al.*) showing binding of AP-1 family members (c-Jun, JunB, JunD) at the *Gli1* promoter. Binding at the *Ifng* promoter serves as a positive control and no binding at the *Gli2* and *Gli3* promoters serve as negative controls.

The three signaling branches of the TCR: NFAT, NF-κB, and AP-1, respectively, can be pharmacologically manipulated. Ionomycin is a Ca^2+^ ionophore that increases cytosolic Ca^2+^ via release of Ca^2+^ from ER stores, leading to activation of NFAT. Phorbol 12-myristate 13-acetate (PMA) activates PKCs and thus NF*κ*B and AP-1 transcription factors. When naïve CD8^+^ T cells or CTLs were treated with both PMA and Ionomycin, to achieve maximum activation of the T cells, *Gli1* RNA was induced 30 and 3-fold, respectively (Fig. 2B). On their own however, only PMA induced significant upregulation of Gli1 in naïve T cells and CTLs, while ionomycin had only a small effect on Gli1 levels at the highest concentration. This suggested that NFAT signaling might be less important in Gli1 induction.

Since mimicking TCR engagement by PMA and Ionomycin is not physiological, we decided to specifically block the three branches of TCR signaling downstream of TCR crosslinking.

The NFAT and NF*κ*B signaling branches can be specifically blocked by transfecting primary CD8^+^ T cells with VIVIT or Dominant Negative (DN)IkB*α*, respectively. VIVIT is a 16mer peptide that selectively disrupts the interface between Calcineurin and NFAT ^16^. As expected, VIVIT-expressing CTLs showed reduced IL-2 and IFN*γ* production (Suppl. Fig. 2A). DN IkB*α* cannot be phosphorylated by IKK*β* and thus NF*κ*B is sequestered in the cytosol ^17^. Indeed, CTLs transfected with DN IkB*α* had reduced nuclear NF*κ*B translocation (Suppl. Fig. 2B). When Gli1 induction was assessed at steady state and after 3 hours of restimulation, VIVIT-and DN IkB*α* -expressing CTLs had no defect in Gli1 induction compared to GFP-transfected controls (Fig. 2C).

The third branch of TCR signaling leading to AP-1 activation can be blocked by U0126, a selective MAP Kinase (MEK1 and MEK2) inhibitor (Suppl. Fig. 2C) ^18^. U0126 treatment did not affect cell viability (Suppl. Fig. 3A), but led to a significant reduction of Gli1 induction after TCR stimulation in CTLs and to a much lesser degree in naïve CD8^+^ T cells (Fig. 2D). MAPK signaling has been shown to be able to activate Gli1 through one of two mechanisms. The first is through direct activation of Gli1 transcription by AP-1 heterodimer which is formed by one Fos and one Jun (c-Jun, JunB or JunD) family member. The second mechanism is via the kinase activity of MAPK which activates an unknown upstream regulator of Gli1 ^11^.

To determine which mode of MAPK signaling is responsible for the Gli1 induction, we reanalysed existing ChIP-Seq datasets generated by *Kurachi et al.* to assess direct AP-1 binding to genomic locus of Gli1 in CTLs ^19^. Since AP-1 has been demonstrated to bind upstream of the *Ifng* locus ^20, 21^, we analysed whether Jun transcription factors bind to this locus in the dataset, and indeed, all three Jun family members bind upstream of the *Ifng* locus (Fig. 2E, top panel). Interestingly, we observed that both c-Jun and JunB also bind upstream of the *Gli1* locus (Fig. 2E). By contrast, no binding was observed upstream of the *Gli2* and the *Gli3* loci, which are both not expressed in CTLs ^10^ (Fig. 2E).

Taken together, this data indicates that of the main downstream signaling arms of the TCR (NFAT, NFkB, AP-1), AP-1 signaling selectively controls Gli1 induction in CTLs, in part by direct binding of AP-1 to the genomic locus of *Gli1*. However, none of the major downstream signaling arms appear to be individually responsible for Gli1 induction in naïve CD8^+^ T cells.

### Ca^2+^ is required for Gli1 induction

We could not clearly associate the induction of Gli1 with one specific branch of TCR signaling in naïve CD8^+^ T cells. While MAPK signaling accounted to some extent for the induction of Gli1 in CTLs, it was still not clear what other pathways drive the induction of Gli1 in CD8^+^ T cells. Ca^2+^ is required for the initiation of canonical Hh signaling at the level of the Ihh:Ptch interaction ^22^ and is an important second messenger in T cells ^23^. We therefore investigated whether Ca^2+^ signaling was important for Gli1 induction. We subjected naïve T cells and CTLs to treatment with a cell-permeable high-affinity Ca^2+^ chelator, BAPTA-AM, prior to TCR stimulation. Ca^2+^ chelation with BAPTA-AM led to complete abrogation of Gli1 induction in naïve T cells and a significant reduction of *Gli1* mRNA and protein induction in CTLs upon TCR stimulation (Fig. 3A). BAPTA-AM can chelate Ca^2+^ in both intra- and extracellular compartments. To determine the relative contribution of each of these pools of Ca^2+^ to Gli1 induction, we repeated the experiments using non-cell-permeable BAPTA or EGTA which both chelate extracellular Ca^2+^. Extracellular Ca^2+^ chelation significantly reduced Gli1 induction in CTLs (Fig. 3B). Thus, Ca^2+^ signaling is essential for Gli1 induction and relies on both extra- and intracellular Ca^2+^ pools.

**Figure 3:**
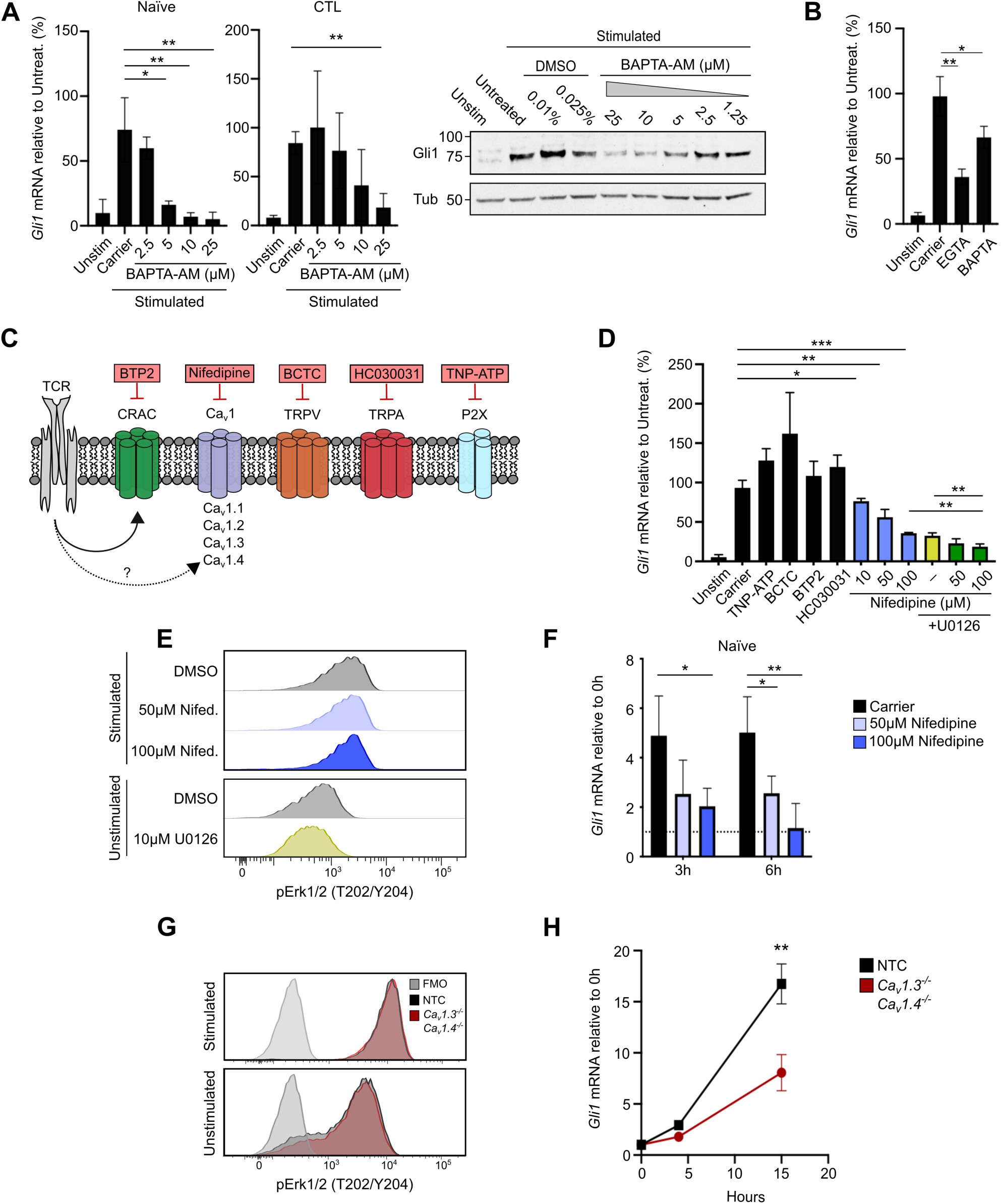
Nifedipine-sensitive Ca^2+^ channels control Gli1 induction in naïve CD8^+^ T cells and CTLs post TCR stimulation. (**A**) Naïve CD8^+^ T cells (left panel) or CTLs (middle panel) were stimulated with plate-bound anti-CD3/CD28 or anti-CD3 antibodies alone for 3 hours in the presence of indicated doses of the cell permeable Ca^2+^ chelator BAPTA-AM or carrier control. RNA was extracted for qRT-PCR analysis. Immunoblot analysis of Gli1 and tubulin of CTLs left unstimulated or restimulated with plate-bound anti-CD3 for 3 hours in the presence of BAPTA-AM or carrier control is shown (right panel). Molecular masses are shown in kilodaltons. n=3 independent experiments. (**B**) CTLs were restimulated for 3h with plate-bound anti-CD3 in the presence of 1.25mM BAPTA, 1.25mM EGTA or carrier control before RNA was extracted for qRT-PCR analysis (right panel). n=3 independent experiments. (**C**) Schematic overview of Ca^2+^ channels with respective antagonists (red) used in this study. (**D**) CTLs were restimulated with plate-bound anti-CD3 in the presence of the indicated inhibitors or carrier control for 15h before RNA was extracted for qRT-PCR analysis. n=3-4 independent experiments. (**E**) CTLs were restimulated with 10µg/ml cross-linked soluble anti-CD3 in the presence of Nifedipine, 10µM U0126 or carrier control. Cells were subsequently prepared for intracellular flow cytometric analysis. n=2 independent experiments. (**F**) Naïve CD8^+^ T cells were stimulated with plate-bound anti-CD3ε in the presence of the indicated doses of Nifedipine or carrier control. n=3 independent experiments. (**G, H**) Activated CD8^+^ T cells were electroporated with RNP complexes at day 2 post stimulation to generate *Cav1.3/1.4^-/-^* (KO) or non-targeting control (NTC) CTLs. CTLs were restimulated on day 8/9 with 10µg/ml cross-linked soluble anti-*CD3ε* for flow cytometric analysis of pErk (**G**) or restimulated with plate-bound anti-CD3 for qRT-PCR analysis (**H**). n=3 independent experiments. (A,B,D,F,H) Data is normalized to *CD3ε* as a reference gene. Similar results were obtained when *Tbp* was used as a reference gene. Error bars indicate SD. p values were calculated using an unpaired two-tailed Student’s t test (A,B,D,F) or two-way ANOVA (H) and * indicates p<0.05, ** p<0.01, and *** p<0.001.

### L-type voltage gated Ca^2+^ channels are critical for Gli1 induction and act independently of MAPK signaling

We next sought to determine whether Ca^2+^ signaling through a specific Ca^2+^ channel is responsible for regulating Gli1 induction. We targeted channels previously shown to be expressed in T cells such as CRAC channels, L-type voltage-gated Ca^2+^ channels (Cav1), TRPV, TRPA and P2X channels with small-molecule inhibitors (Fig. 3C) ^23^. Treatment of CTLs with small molecule inhibitors at doses previously published in the literature had no effect on cell viability during the course of the experiment (Suppl. Fig. 3A). Interestingly, inhibition of CRAC channels showed no effect on Gli1 induction and neither did inhibition of TRPV, TRPA and P2X channels.

Only inhibition of Cav1 channels with Nifedipine showed a dose-dependent reduction of Gli1 induction in CTLs (Fig. 3D). Cav1 channels have been shown to play a critical role in CD8^+^ T cell development ^24^. Evidence in the literature has shown that constitutive knockout of *CACNA1F* (encoding Cav1.4) during T cell development results in impaired MAPK signaling strength ^12^. We hence sought to determine whether the effect of Nifedipine on Gli1 induction in CTLs was through inhibition of MAPK signaling or through an independent mechanism. Inhibition of both, MAPK and Cav1 channels, showed an additive decrease in Gli1 induction (Fig. 3D) but importantly pErk levels were unaffected by treatment with Nifedipine (Fig. 3E), indicating that Cav1 channels were regulating Gli1 induction through a mechanism distinct from MAPK signaling. Given that MAPK signaling is not a major driver of Gli1 induction in naïve CD8^+^ T cells (Fig. 2D) we sought to investigate whether Cav1 channels might be the main source of Gli1 induction. Nifedipine treatment of naïve CD8^+^ T cells resulted in complete abrogation of Gli1 induction post TCR stimulation (Fig. 3F).

We wanted to confirm this finding by genetic ablation of Cav1 channels and performed CRISPR knockouts of each of the known Cav1 α1 subunit coding genes (*CACNA1S/C/D/F*) in primary murine CD8^+^ T cells. Knockout of individual genes did not show a significant decrease in Gli1 induction (Suppl. Fig. 4A) indicating that there may be a degree of redundancy between individual Cav1 family channels. Cav1.3 and 1.4 KO led to the most sustained reduction in Gli1 induction. Thus, we next performed a double knockout of *CACNA1D and CACNA1F*, which encode Cav1.3 and Cav1.4, respectively. The double knockouts showed high knockout efficiency (Suppl. Fig. 4B) and normal cell proliferation (Suppl. Fig. 4C). T cells double knockout for Cav1.3 and Cav1.4 had intact MAPK signaling (Fig. 3G) but a striking inability to induce Gli1 upon stimulation (Fig. 3H).

Thus, we have shown that Cav1 channels are the main inducers of Gli1 in naïve CD8^+^ T cells and in CTLs act together with MAPK signaling to induce Gli1.

### Cav channels localize to the plasma membrane and intracellular vesicles in human CD8^+^ T cells and are important for killing

To interrogate whether Cav1 channels are also expressed in human CD8^+^ T cells we profiled the expression of Cav1 α1 subunit coding genes (CACNA1S/C/D/F corresponding to Cav1.1, Cav1.2, Cav1.3, and Cav1.4, respectively) in naïve CD8^+^ T cells from multiple healthy donors (Fig. 4A). While Cav1.1 and Cav1.2 were not expressed in human CD8^+^ T cells, we readily detected expression of Cav1.3 and Cav1.4, the two subunits responsible for Gli1 induction in murine T cells with Cav1.4 being 50-fold higher expressed than Cav1.3.

**Figure 4:**
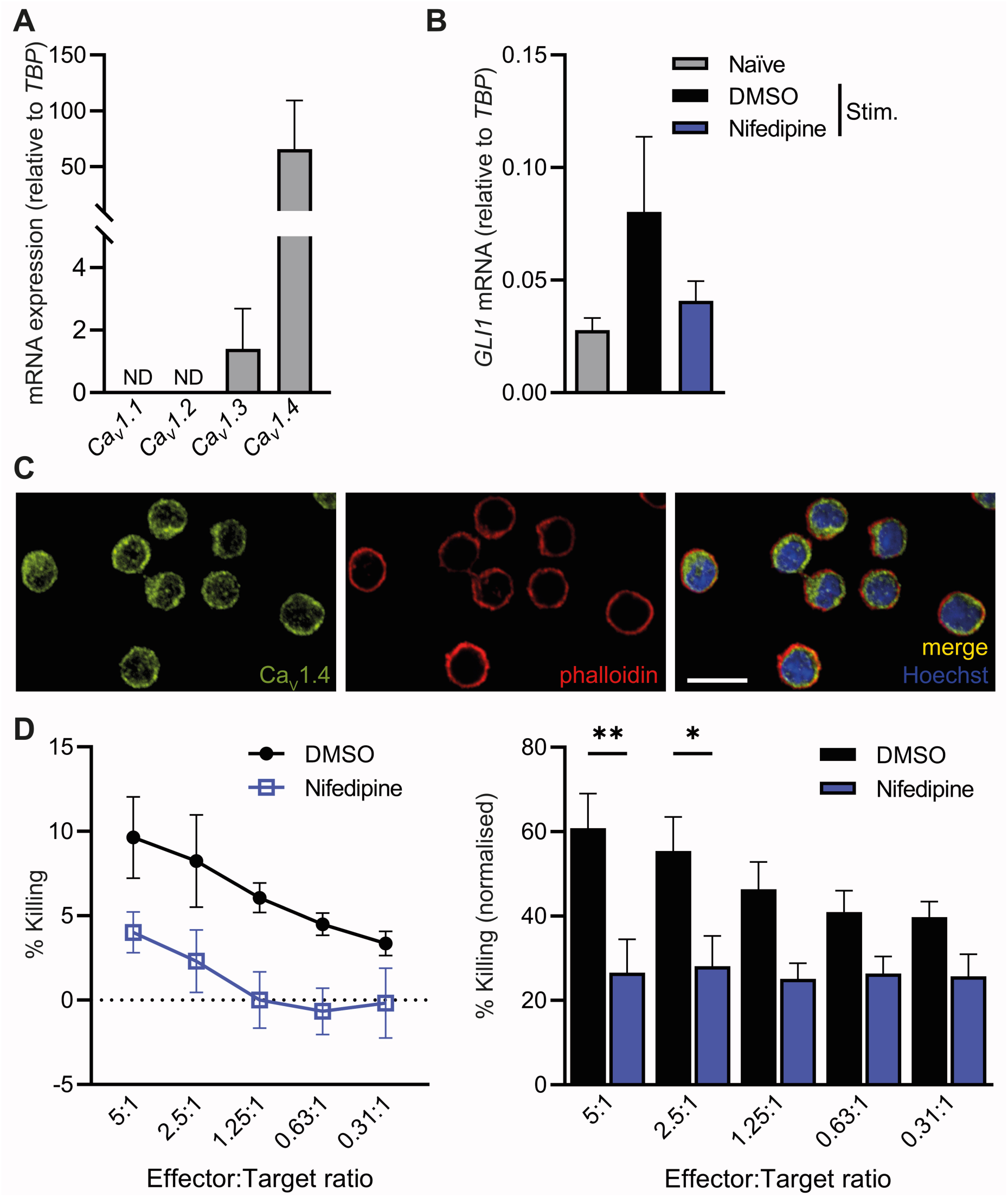
Cav1 family channels are responsible for Gli1 induction and control killing in human CTLs. Naïve human CD8^+^ T cells were isolated from peripheral blood of healthy donors. (**A**) Cells were processed for qRT-PCR analysis of human Cav1.1 (*CACNA1S*), Cav1.2 (*CACNA1C*), Cav1.3 (*CACNA1D*) and Cav1.4 (*CACNA1F*) channel expression. Values were normalized to *TBP* as a reference gene. ND: Not Detected. Error bars indicate SEM. n=7 donors. (**B**) RNA was extracted before and after stimulation with human T-activator CD3/CD28 beads for 2 days in the presence of 100 µM nifedipine or DMSO (carrier control). Error bars indicate SEM. n=11 donors. (**C**) T cells were stimulated with human T-activator CD3/CD28 beads for 3 days. On d11, CTLs were stained with antibodies against CaV1.4, phalloidin and Hoechst. Scale bar=10 µm. n=8 donors, representative donor shown. (**D**) T cells were stimulated with human T-activator CD3/CD28 beads for 3 days and again restimulated with T-activator CD3/CD28 beads on d10 for 3 days. On d14-d15, CD8^+^ T cells were co-cultured with P815 target cells at indicated effector to target ratios and subjected to an LDH cytotoxicity assay in the presence of 100µM Nifedipine or DMSO (carrier control). Left panel: Error bars indicate SD, representative donor shown. Right panel: Error bars indicate SEM, n=6 donors. p values were calculated using two-way ANOVA. * indicates p<0.05, and ** p<0.01.

Next, we wanted to know whether the Cav1 channel inhibitor Nifedipine could block GLI1 induction in human CD8^+^ T cells as it did in murine cells (Fig. 3F). The concentration of Nifedipine used had no effect on cell viability (Suppl. Fig. 3B) but induction of GLI1 was ablated in Nifedipine-treated human CD8^+^ T cells (Fig. 4B).

Cav1 channels have been hypothesized to localize to the plasma membrane in T cells ^12^ but have never been localized by immunofluorescence. Using an antibody against Cav1.4 ^25^, we stained human CD8^+^ T cells from different healthy donors and consistently observed specific staining at the plasma membrane as well as on intracellular vesicles in all donors (Fig. 4C).

To investigate whether Cav1 channels are implicated in the ability of human CD8^+^ T cells to kill tumor targets, we treated human CTLs from various donors with Nifedipine or carrier control before subjecting them to a killing assay. While carrier-treated human CTLs were able to kill tumor targets, Nifedipine-treated CTLs from all donors tested showed a marked decrease in killing ability (Fig. 4D).

Taken together, we have shown that, similarly to murine CD8^+^ T cells, human CD8^+^ T cells express Cav1 channels and depend on them for GLI1 upregulation and consequent target cell killing.

### L-type voltage gated Ca^2+^ channels regulate T cell killing in a Gli1-dependent manner

To definitively answer whether the effect of Cav1 channel blockade via Nifedipine on CTL killing is achieved through Gli1 or an independent mechanism, we treated *Gli1* WT and *Gli1* KO CTLs with Nifedipine or carrier control (Fig. 5A). We find that Nifedipine treatment leads to a defect in killing in *Gli1* WT CTLs, which we have shown to be independent of effects on cell viability or MAPK signaling (Fig. 3E, Suppl. Fig. 3A). This indicates that Cav1 channels are important regulators of CTL killing. Treatment of *Gli1* KO CTLs with Nifedipine showed no additive defect in killing compared to carrier control (Fig. 5A), indicating that the presence of Gli1 is required for Nifedipine to exert its effect on CTL killing.

**Figure 5:**
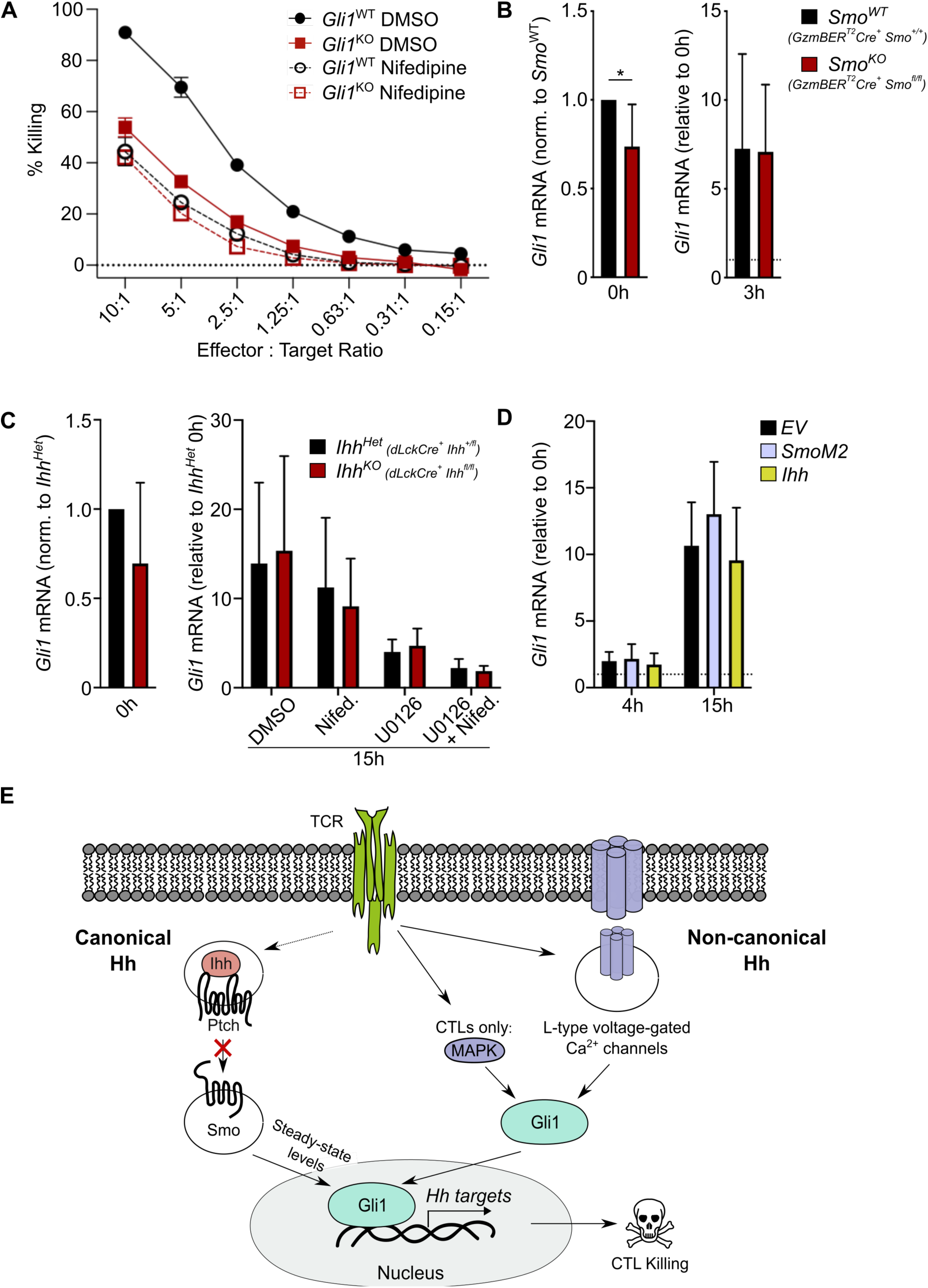
Cav1 family channels control CTL killing in a Gli1-dependent manner and function independently of canonical Hh signaling. (**A**) CTLs were generated from *Gli1^WT^*, *Gli1^Het^* and *Gli1^KO^* mice. On day 7, cells were co-cultured with either peptide-pulsed EL-4 cells (OTI) or anti-CD3 antibody-coated P815 target cells at the indicated effector to target ratios and subjected to an LDH cytotoxicity assay in the presence of 100μM Nifedipine or carrier control. Error bars indicate SD. Representative of n=4 independent experiments. (**B**) CTL were generated from *GzmBERTCre^+^ Smo^+/+^* (*Smo^WT^*) or *GzmBERTCre^+^ Smo^fl/fl^* (*Smo^KO^*) mice. Cells were treated with 4-OH-Tamoxifen for the duration of the *in vitro* experiments. CTLs were restimulated for 3h on day 8/9 for qRT-PCR analysis. n=5 mice per condition, 5 independent experiments. (**C**) CD8^+^ T cells were isolated from *dLckCre^+^ Ihh^fl/+^* (*Ihh^Het^*) or *dLckCre^+^ Ihh^fl/fl^* (*Ihh^KO^*) mice. CTLs were restimulated for 15h on day 8/9 in the presence of 100µM Nifedipine and/or 10µM U0126 or carrier control for qRT-PCR analysis. n=5 (naïve) and n=3 (restimulation) independent experiments. (**D**) CD8^+^ T cells were retrovirally transduced with pMig constructs encoding empty vector (EV), SmoM2 or Ihh, respectively. Sorted, transduced CTLs were restimulated on day 8/9 for qRT-PCR analysis. n=3 mice per condition, 3 independent experiments. (B-D) Data is normalized to *CD3ε* as a reference gene. Similar results were obtained when *Tbp* was used as a reference gene. Error bars indicate SD. No statistically significant differences were observed. (**E**) Model of TCR-induced Gli1 induction: two arms of Hedgehog (Hh) signaling are induced post-TCR stimulation. Canonical Hh signaling is initiated by binding of intracellular Ihh to Ptch, which results in activation of Smo. Smo is the key signal transducer of canonical Hh signalling and was previously shown to regulate steady-state Gli1 protein expression (*de la Roche et al*). TCR stimulation also activates Cav channels and we show here that these Cav channels are critical to induce Gli1 transcription either independently (naïve CD8^+^ T cells) or in conjunction with MAPK signalling (CTLs). The TCR-induced Gli1 activation is essential for efficient CTL killing.

Thus, we have shown that L-type voltage gated Ca^2+^ channels are critical for CTL killing and that the mechanism for this killing phenotype is by regulating the induction of Gli1.

### Gli1 induction upon TCR signaling is independent of canonical Hh signaling via Ihh and Smo

Given the role of Smo in maintaining Gli1 steady-state protein expression ^10^ we asked whether canonical Hh signaling was needed for the induction of Gli1 after TCR stimulation. To investigate this, we generated two different conditional knockout lines (Suppl. Fig. 5A,B). For the first mouse line, we crossed *GzmB ER^T2^Cre* mice ^26^ to *Smo^f/f^* mice, to generate mice in which Cre is only active in mature CTLs. The second mouse line was generated by crossing *dLckCre* mice to *Ihh^f/f^* mice, in which the Cre is active only after T cell development has been completed in the thymus. Ihh is the only Hh ligand expressed by CD8^+^ T cells and Hh signaling has been suggested to be entirely intracellular in CD8^+^ T cells ^10^. We therefore decided to knock out endogenous Ihh in CD8^+^ T cells as well as Smo the key signal transducer of canonical Hh signaling. Importantly, the peripheral T cell compartment in conditional *Ihh* KO mice and inducible *Smo* KO mice is normal (Suppl. Fig. 5A,B). Interestingly, CD8^+^ T cells from *Smo^f/f^* and *Ihh^f/f^* mice showed a modest reduction of *Gli1* mRNA at steady state but upregulated *Gli1* mRNA to the same extent as WT cells upon TCR stimulation (Fig. 5B,C), indicating that Gli1 induction upon TCR signaling is not mediated by canonical Hh signaling.

Ca^2+^ is required for the binding of Ihh to Ptch ^22^, thought to initiate canonical Hh signaling in CD8^+^ T cells. To confirm that canonical Hh signaling is indeed independent of our newly identified Cav1-regulated Gli1 induction, we treated CTLs from *Ihh^+/fl^* and *Ihh^fl/fl^* mice with Nifedipine and U0126. We observed the same levels of inhibition of Gli1 induction in both conditions (Fig. 5C), suggesting that the MAPK-Cav-Gli1 axis is independent of canonical Hh signaling.

We also considered whether overexpression of either Ihh or Smo would further increase Gli1 induction. For this we transduced primary CD8^+^ T cells with retroviruses encoding Ihh or a constitutively active form of Smo (SmoM2) ^27^, resulting in 80000- and 300-fold overexpression, respectively (Suppl. Fig. 5C). Strikingly, we did not observe any further increase in the induction of Gli1 as compared to empty vector-transduced control CD8^+^ T cells. (Fig. 5D).

Thus, canonical Hh signaling via Ihh and Smo is not responsible for the induction of Gli1 downstream of the TCR, and is independent of the MAPK and L-type voltage-gated Ca^2+^ channel-induced Gli1 activation.

## DISCUSSION

In this manuscript we have uncovered a novel non-canonical Hh signaling pathway whereby a Cav1 channel-mediated Ca^2+^ flux can induce Gli1. We show that this mechanism of Gli1 induction is functionally critical for CTL killing, providing a direct mechanistic link between Cav1 channels and CTL killing (Fig. 5E).

There is currently great scientific interest to unmask the roles of Cav channels in T cell biology. Constitutive knockout of Cav1.4 or the Cav channel regulatory *β*3 subunit leads to signaling defects in thymocytes and apoptosis of peripheral T cells, respectively ^13^. This makes it challenging to draw conclusions about the role of Cav channels in mature peripheral T cells. Here we use CRISPR technology to functionally ablate Cav channels in primary mature T cells only.

All four family members of the L-type Ca^2+^ channels (Cav1.1, Cav1.2, Cav1.3, and Cav1.4) are expressed in murine T cells ^24^ and we find that only Cav1.4 and to a lesser extent Cav1.3 are expressed in human CD8^+^ T cells. We show that Cav1.3 and Cav1.4 (and to a lesser extent Cav1.1) are involved in the induction of Gli1 in CD8^+^ T cells. Apart from the localization of Cav1.4 to lipid rafts little is known about the cellular localization of Cav channels ^12^. Here we find that Cav1.4 is expressed at the plasma membrane and on intracellular vesicles in primary human CD8^+^ T cells.

There is evidence from the literature that in neuronal cells canonical Hh signaling can induce Ca^2+^ spikes via Cav and TRPC1 channels ^7^. Here we show that the “reverse” is true in T cells, where we show that Ca^2+^ signaling via Cav channels induces Hh signaling. Our work uncovers Ca^2+^ signaling via Cav channels as a novel mechanism of non-canonical, Ptch/Smo-independent Hh signaling.

The literature supports the notion that our proposed Cav-Gli axis may be active in other cells. Humans with mutations in Cav channels phenotypically display heart arrhythmias and signs of autistic spectrum disorder (consistent with the critical roles of Cav channels in the heart and brain) as well as signs of syndactyly ^28^, a phenotype which has been robustly linked to inactivating mutations in the Hh pathway ^29^. Furthermore, studies in mice showed that knockdown of Cav channels leads to abnormalities in skeletal development ^30^, highly reminiscent of the defects observed in *Ihh* KO mice ^31^. These studies suggest that loss-of-function phenotypes of Cav channels phenotypically mimic known Hh loss-of-function phenotypes, consistent with the hypothesis that a Cav-induced mechanism of Gli activation is operational in other cell types.

Interestingly, we uncover that in CTLs MAPK activation downstream of the TCR critically contributes to Gli1 transcription which has been shown in other cell types and cancers ^11^. Of interest is the difference of Gli1 regulation between naïve CD8^+^ T cells, that seem to solely rely on Cav signaling, and CTLs, that also employ MAPK signaling. A possible explanation for this could be that naïve CD8^+^ T cells have much longer contact times with an APC and form more stable synapses *in vivo* when they are stimulated than CTLs, which have shorter contact times with their target cells and have been shown to often require multiple rounds of contact *in vivo* to efficiently kill target cells ^32^. Therefore, CTLs may have evolved two pathways to ensure appropriate levels of Gli1 activation.

We show that Gli1 activation by Cav channels is important for killing in murine and human CD8^+^ T cells (Fig. 5A, Fig. 4E) which is a rapidly-induced process with killing observed within a few hours *in vitro*. This is consistent with findings in the literature that Gli1 activation can be rapidly induced ^33^, but the question remains how the transcription factor Gli1 can exert its effect on killing in such a short timeframe. Given that Gli1 can regulate the cytoskeleton ^10^, we hypothesize that Gli1-induced transcription in CTLs upon target recognition is required for cytoskeletal rearrangement and immune synapse formation upon successive target cell contacts and serial killing.

Regarding the activation of Cav channels in T cells, it is tempting to speculate that this is PKC-driven because of two observations. First, Cav channels in T cells lack the ability to respond to depolarization and instead become activated upon TCR engagement ^34^. And second, PKC is a well-known activator of Cav channels in excitable cell types and has been hypothesized as the link between TCR and Cav channel activation in T cells ^35, 36^. This would explain why we find that PMA (a PKC agonist) is able to drive Gli1 induction (Fig. 2B), particularly in naïve CD8^+^ T cells where this effect cannot simply be explained by PKC-related induction of MAPK signaling (which has no effect on Gli1 induction). The requirement for PKC for Cav activation and the different nature of the Ca^2+^ flux induced by ionomycin might also explain why ionomycin administration alone cannot induce Gli1.

The precise mechanism by which Cav1 channel activation results in Gli1 induction is not known at this point. One potential mechanism is via Ulk family proteins that can be activated by Ca^2+^ and have been shown to activate Gli1 via proteolytic cleavage in a very short timeframe ^33, 37^. And indeed we find Ulk3 expressed in CD8^+^ T cells (Suppl. Fig. 5D).

Canonical Hh signaling is active for long periods of time, is initiated by extracellular ligands, and computes a cell fate choice ^3^. By contrast, CTL killing must happen much more rapidly and should not rely on extracellular ligand gradients. Canonical Hh signaling is a more ancient signaling pathway than TCR signaling ^2^. The pathway might have come under evolutionary pressure for the induction of Gli1 to be more rapid and isolated from extracellular signals resulting in direct activation by the TCR.

Taken together, we show that Gli1 is critical for CD8^+^ T cell killing of tumor cells and that Gli1 activation is intricately coupled to CD8^+^ T cell activation downstream of the TCR via Cav channels. Our findings have clinical implications. Gli inhibitor Arsenic Trioxide (ATO) is currently in clinical trials for the treatment of various cancers including advanced Basal Cell Carcinoma and Acute Myeloid Leukemia ^38^. Our data suggests that ATO may inhibit the CD8^+^ anti-tumor response and might explain why Gli1 inhibitors have had little success in the clinic so far.

## MATERIALS AND METHODS

### Mice

RAG2KO were a generous gift from Suzanne Turner (University of Cambridge) and OTI mice were purchased from the Jackson Laboratory (C57BL/6-Tg(TcraTcrb)1100Mjb/j, Stock no. 003831). OTI RAG2KO mice were generated from these. *Gli1-eGFP* mice were a generous gift from Alexandra Joyner (Sloan Kettering Institute)^39^ and were backcrossed onto the C57BL/6J background (The Jackson Laboratory, Bar Harbor, ME) for more than 11 generations. *dLckCre* and *ROSA26tdTom* mice were a generous gift from Randall Johnson and Douglas Winton (University of Cambridge), respectively. GzmBER^T2^*Cre/ROSA26EYFP* mice were a generous gift from D. T. Fearon (Cold Spring Harbor Laboratory)^26^ and were backcrossed onto the C57BL/6J background (The Jackson Laboratory, Bar Harbor, ME) for more than 11 generations. *Smo^f/f^* (Smo^tm2Amc^/J, Stock no. 004526) and *Ihh^f/+^* (*Ihh^tm1Blan^*/J, Stock no. 024327) mice were purchased from The Jackson Laboratory. *Smo^fl/fl^* mice were back-crossed to the C57BL/6J background (Charles Rivers Inc., UK) for more than 10 generations and crossed to GzmBER^T2^*Cre/ROSA26EYFP* mice to generate *GzmBER^T2^Cre/ROSAEYFP/Smo^fl/fl^* mice. *Ihh^fl/fl^* mice were crossed onto *dLckCre/ROSA26tdTom* mice.

Mice were maintained at the CRUK Cambridge Institute/ University of Cambridge, genotyped using Transnetyx, and used at 6-8 weeks of age. All housing and procedures were performed in strict accordance with the United Kingdom Home Office Regulations.

### Cell Culture

Purified naïve murine CD8^+^ T cells were cultured at 1x10^6^ cells/ml in complete RPMI media: RPMI (Gibco, Cat no. 21875034) supplemented with 5% heat-inactivated batch-tested FCS (Biosera, Cat no. 1001-500ml), 50µM β-Mercaptoethanol (50mM, Gibco, Cat no. 31350-010), 100U/ml Penicillin/Streptomycin (10’000 U/ml, Gibco, Cat no.15140-122), 1µM Sodium Pyruvate (Gibco, Cat no. 11360039), 10µM HEPES (Sigma, Cat no. H0887) and 10ng/ml murine IL-2 (Peprotech, Cat no. 212-12). Human CD8^+^ T cells were cultured in ImmunoCult cell culture media (Stemcell Technologies, Cat no. 10981) supplemented with 35 ng/ml human IL-2 (Peprotech, Cat no. 200-02).

HEK 293T cells and P815 cells were cultured in DMEM (Gibco, Cat no. 21885-025) supplemented with 10% heat-inactivated FCS (Biosera, Cat no. 1001-500ml). EL-4 cells were cultured in RPMI supplemented with 10% heat-inactivated FCS.

All cells were grown in a humidified incubator at 37°C and 5% CO2 and cell lines tested mycoplasma negative (MycoProbe*^®^* Mycoplasma Detection Kit, R&D systems).

### CD8^+^ T cell isolation and activation

#### Mouse

Spleens were harvested and cell suspensions made. Naïve murine CD8^+^ T cells were isolated using negative selection (Naïve CD8 T Cell Isolation Kit, Cat no. 130-096-543, Miltenyi Biotec) according to the manufacturer’s instructions. The purity of the sorted populations was above 95%. CD8^+^ T cells were stimulated with platebound anti-CD3*ε* (1µg/ml, eBioscience, Cat no. 16-0033-86) and anti-CD28 (2µg/ml, eBioscience, Cat no. 16-0281-86) antibody.

For CTL generation, a splenocyte suspension from one OT-I Rag2^-/-^ spleen was stimulated with 10nM Ova257-264 peptide (SIINFEKL, AnaSpec, Cat no. 60193-5-ANA) for 48h and cultured for up to 10 days. Cells were restimulated with plate-bound anti-CD3*ε* (2.5μg/ml).

#### Human

Buffy Coats from healthy donors were acquired from NHS Blood and Transplant (Cambridge) under appropriate ethics (Research into Altered Lymphocyte Function in Health and Disease, REC reference: 17/YH/0304). PBMCs were isolated using SepMate PBMC Isolation Tubes (Stemcell Technologies, Cat no. 86450) according to the manufacturer’s instructions and naïve CD8^+^ cells were obtained using the MojoSort Human CD8^+^ Naïve T Cell Isolation Kit (Biolegend, Cat no.480045). Purity was consistently over >75% CD8^+^ with no contamination of CD4+ T cells and >80% of CD8+ T cells being naive. Cells were >95% CD8^+^ on day 1of stimulation. Naïve CD8^+^ cells were plated with Human T-Activator CD3/CD28 Dynabeads (Thermofisher, Cat no.111.31D) at 25 µl of Dynabeads per 1x10^6^ CD8^+^ cells in complete ImmunoCult cell culture media (see above). On day 3, the beads were magnetically removed. When indicated, the cells were re-stimulated on d10 with 12.5 µl of Dynabeads per 1x10^6^ CD8^+^ cells for three additional days. On d14, the beads were removed.

### CRISPR knockout of primary CD8^+^ T cells

Ribonucleoprotein (RNP) complexes were assembled following the manufacturer’s instructions using Alt-R^®^ CRISPR-Cas9 crRNA (IDT) and tracrRNA (IDT, Cat no. 1072534). CD8^+^ T cells were isolated from OT-I Rag2^-/-^ spleens and stimulated with plate-bound anti-CD3*ε*/CD28 antibodies in complete RPMI media. After 48h of stimulation, cells were washed twice in pre-warmed PBS (Gibco) prior to resuspension in 80µl Buffer T (10^6^ cells/electroporation reaction). The suspension was briefly mixed with the RNP complex solution prior to electroporation with the Neon™ electroporation system in 100µl electroporation tips (Thermo, Cat no. MPK10096) with three pulses of 1600V each with a pulse width of 10ms. Cells were left to recover in complete RPMI in the absence of antibiotics for 20min. Cells were then centrifuged at 1500rpm for 5min and returned into culture in complete RPMI media (without antibiotics). Antibiotics were re-added after six hours at the indicated concentration.

### Genome editing efficiency assay

Genomic DNA was extracted from T cells using a DNeasy Blood & Tissue kit (Qiagen, Cat no. 69506). PCR amplification of the region of gDNA containing the CRISPR editing locus was performed using the SequalPrep Long PCR Kit with dNTPs (Invitrogen, Cat no. A10498) according to the manufacturer’s instructions. Primers (Sigma) used were as follows:

A T7 endonuclease I mismatch cleavage assay was performed using the Alt-R® Genome Editing Detection Kit (IDT, Cat no. 1075932) per the manufacturer’s instructions. Quantification of cleaved bands was performed on a capillary electrophoresis system (4200 Tapestation, Agilent) using D5000 screentape (Agilent, Cat no. 5067-5588/9).

### *In vitro* killing assay

CD8^+^ T cell cytotoxicity was assessed using a Cyto Tox 96® Non-radioactive Cytotoxicity Assay kit (Promega, Cat no. G1780) according to the manufacturer’s instructions. Murine and human CTLs were generated as previously described and used on day 6/7 or day 14/15 post-stimulation, respectively. EL-4 cells were used as target cells for murine OTI CTLs and pulsed for 1h at 37°C with 1µM SIINFEKL. P815 cells were used as target cells for human CTLs and pulsed for 1h at 37°C with 1µg/ml anti-CD3 antibody (clone UCHT1, Biolegend, Cat no. 300438). They were washed and resuspended at 1x10^5^ cells/ml in killing assay media (RPMI without phenol red + 2% FCS) in a round-bottom 96 well plate.

T cells were washed, resuspended in killing assay media and plated at the indicated effector:target ratios. Plates were incubated at 37°C prior to collection of supernatant at the designated timepoints. Absorbance was measured at 490nm using a CLARIOstar microplate reader (BMG Labtech).

### Flow cytometry and FACS

#### Surface staining

Cells were stained with fixable viability dye eFluor780 (eBioscience, Cat no. 65-0865-18) or DAPI (Thermo Fisher, Cat no. D9542, 1:3000 in PBS), washed, and incubated with Fc block (1:100; Biolegend TruStain fcX, cat no. 101320) and fluorophore-conjugated antibodies at the appropriate concentrations (Table 3) for 20min at 4°C.

**Table 1.**
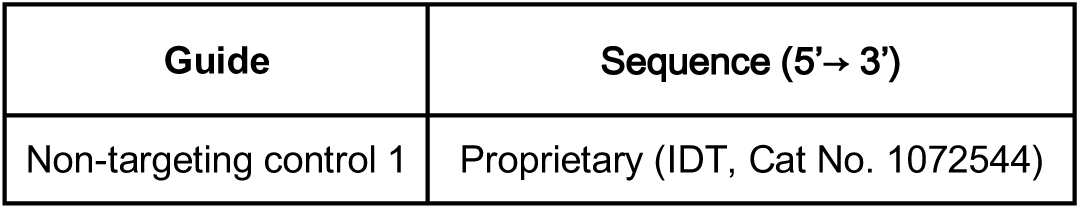
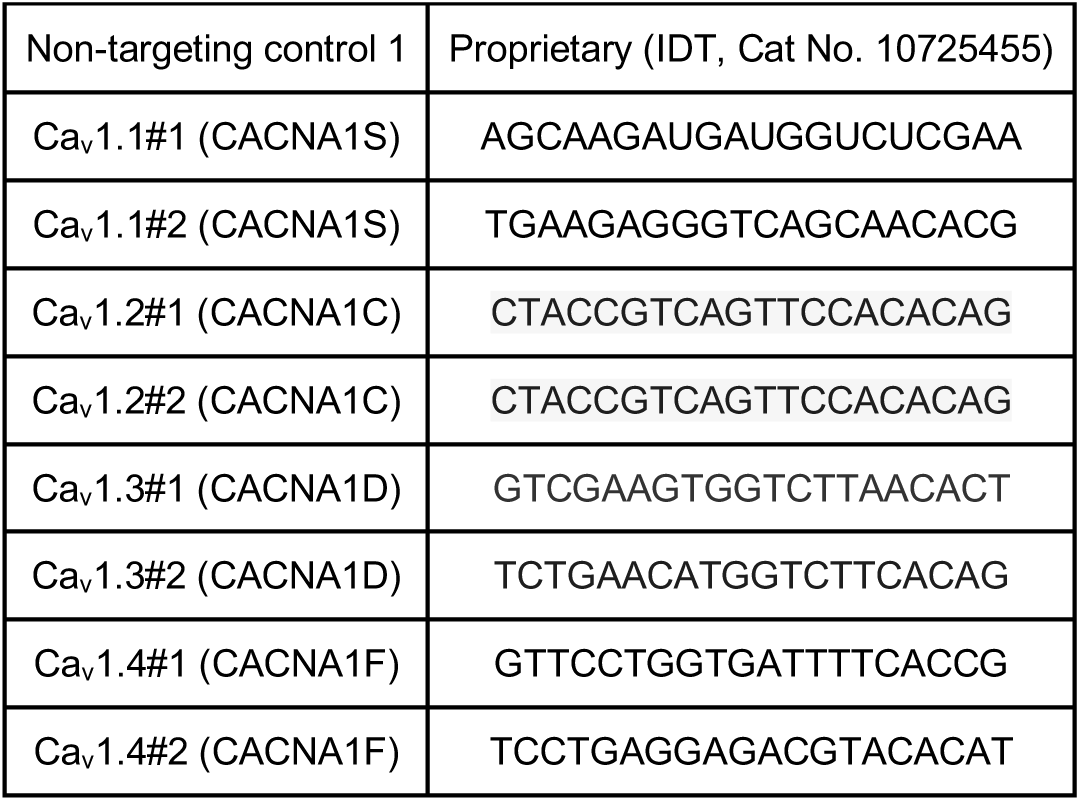
List of crRNA used for CRISPR in CD8+ T cells

**Table 2:**
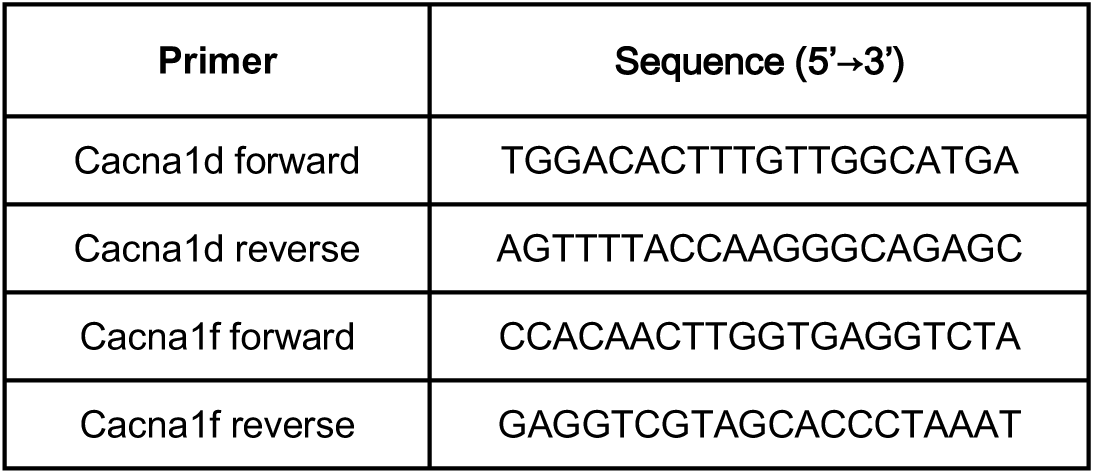
PCR Primers used for genome editing efficiency assays

**Table 3:** List of antibodies used for flow cytometry

| Target | Clone | Reactivity | Fluorochrome | Dilution | Cat no. | Supplier |
| --- | --- | --- | --- | --- | --- | --- |
| CCR7 | G043H7 | Human | PE/Cy7 | 1:50 | 353226 | Biolegend |
| CD3ε | 145-2C11 | Mouse | BUV395 | 1:50 | 583565 | BD |
| CD3ε | UCHT1 | Human | BUV395 | 1:100 | 563556 | BD |
| CD4 | RM4-5 | Mouse | BV605 | 1:200 | 100548 | Biolegend |
| CD4 | RM4-5 | Mouse | APC | 1:200 | 100516 | Biolegend |
| CD4 | RM4-5 | Mouse | FITC | 1:200 | 100510 | Biolegend |
| CD4 | RM4-5 | Mouse | PE | 1:200 | 100512 | Biolegend |
| CD4 | RPA-T4 | Human | BV605 | 1:100 | 300556 | Biolegend |
| CD8a | 53-6.7 | Mouse | BV605 | 1:200 | 100744 | Biolegend |
| CD8a | 53-6.7 | Mouse | APC | 1:200 | 100712 | Biolegend |
| CD8a | 53-6.7 | Mouse | FITC | 1:200 | 100706 | Biolegend |
| CD8a | 53-6.7 | Mouse | PE | 1:200 | 100707 | Biolegend |
| CD8 | RPA-T8 | Human | BV711 | 1:100 | 301044 | Biolegend |
| CD27 | LG.3A10 | Human | PerCP/Cy5.5 | 1:50 | 124214 | Biolegend |
| CD28 | CD28.2 | Human | BV785 | 1:50 | 302950 | Biolegend |
| CD44 | IM7 | Mouse/Human | BV785 | 1:400 | 103059 | Biolegend |
| CD44 | IM7 | Mouse/Human | BV711 | 1:400 | 103057 | Biolegend |
| CD44 | IM7 | Mouse/Human | PE/Cy7 | 1:400 | 103030 | Biolegend |
| CD45 | 30-F11 | Mouse | PE/Cy7 | 1:200 | 103114 | Biolegend |
| CD45 | HI30 | Human | FITC | 1:100 | 304006 | Biolegend |
| CD45RO | UCHL-1 | Human | APC | 1:100 | 304210 | Biolegend |
| CD62L | MEL-14 | Mouse | BV421 | 1:200 | 104436 | Biolegend |
| CD62L | MEL-14 | Mouse | BV510 | 1:200 | 104441 | Biolegend |
| CD90.1 | HIS51 | Mouse/Rat | APC | 1:200 | 17-0900-82 | BD |
| CD90.1 | HIS51 | Mouse/Rat | PE | 1:200 | 12-0900-83 | Thermo |
| CD95 | DX2 | Human | BV421 | 1:50 | 305624 | Biolegend |
| IFNγ | XMG1.2 | Mouse | BV711 | 1:100 | 505835 | Biolegend |
| IL-2 | JES6-5H4 | Mouse | PE | 1:100 | 503808 | Biolegend |
| TCRβ | H57-597 | Mouse | BV711 | 1:200 | 109243 | Biolegend |
| TCR$\gamma\delta$ | GL3 | Mouse | Alexa Fluor 488 | 1:200 | 118128 | Biolegend |
| pErk | 197G2 | Mouse/Human | none | 1:200 | 4377 | Cell Signaling |
| p65 | 532301 | Mouse/Human | APC | 1:20 | IC5078A | R&D Systems |
| Rabbit IgG | Polyclonal | Rabbit | Alexa Fluor 647 | 1:500 | A-21245 | Thermo Fisher |

#### Intracellular staining

Following surface staining, cells were fixed with BD Cytofix/Cytoperm Plus Fixation Buffer (BD Biosciences, Cat no. 554715) as per manufacturer’s instructions and stained with fluorophore-conjugated antibodies at the appropriate concentrations (Table 3) for 30min at 4°C. Prior to analysis, cells were washed once in permeabilization buffer and once in FACS buffer.

#### pErk staining

Naïve CD8^+^ T cells or CTLs were isolated/maintained as described previously. Cells were pre-incubated for 30min at 37°C with 10µM MEK1/2 inhibitor U0126 (Cambridge Biosciences, Cat no. SM106-5) or carrier control (DMSO). Cells were stimulated by crosslinking the TCR using 10µg/ml soluble anti-CD3*ε* and 5µg/ml soluble anti-CD28 with the addition of goat-anti-hamster IgG (1:300) for 2min in a 37°C waterbath. Cells were fixed in 2% PFA (EMS, Cat no. 15710-S), washed and incubated in BD Phosflow Lyse/Fix Buffer (BD, Cat no. 558049) for 10min at 37°C. Cells were washed with BD Phosflow Perm/Wash I Buffer (BD, Cat no. 557885) and stained with primary anti-pErk antibody(30min, 4°C) and then secondary anti-Rabbit Alexa Fluor 647 and anti-CD8a PE antibodies at the indicated concentrations (Table 3) (20min, 4°C).

Flow cytometric analyses were conducted on a BD LSRII or BD LSR Fortessa cell analyser, and data was analysed with FlowJo software (Tree Star Inc., version 10.4).

### *In vitro* tamoxifen treatment

*In vitro* tamoxifen was given as 4-hydroxytamoxifen (4-OH-Tamoxifen or OHT) (TOCRIS, Cat no. 3412) solubilized in DMSO (Life Technologies, Cat no. D12345) in order to activate CreER^T2^ recombinase *in vitro*. Stock solutions were made at 100mM and kept at -20°C protected from light. Cells were treated at a concentration of 300nM, diluted in complete RPMI (Gibco), and fresh 4-OHT was added daily for the duration of culture. Control-treated cells received DMSO in complete RPMI.

### Small molecule treatment of CD8^+^ T cells

For small molecule treatment studies, naïve CD8^+^ T cells or CTLs were incubated for 1h (2h in the case of BTP2 –^40^) prior to (re-)stimulation. Doses of Ca^2+^ channel blockers were chosen based on doses established for use in T cells in the literature (see Table 4). All inhibitors were reconstituted in DMSO (Life Technologies, Cat no. D12345) according to the manufacturer’s instructions and a carrier control was included for all experiments. The small molecules used were as follows:

**Table 4:** Overview of small molecules used

| Inhibitor | Dose(s) | Stock concentration | Manufacturer | Cat no. |
| --- | --- | --- | --- | --- |
| PMA | 1, 5, 10, 50ng/ml | 1mg/ml | Sigma | P1585 |
| Ionomycin | 10, 50, 100, 1000ng/ml | 1mg/ml | Sigma | I9657 |
| U0126<sup>18</sup> | 10μM | Reconstituted freshly | Cambridge Bioscience | SM106-5 |
| BAPTA-AM<sup>41</sup> | 1.25, 2.5, 5, 10, 25μM | Reconstituted freshly | Thermo Fisher | B1205 |
| TNP-ATP<sup>42</sup> | 30μM | 10mM | Tocris | 2464/5 |
| BCTC<sup>43</sup> | 1μM | 10mM | Tocris | 3875/10 |
| BTP2 (YM-58483)<sup>40</sup> | 200nM | 5mM | Tocris | 3939/10 |
| HC030031<sup>44</sup> | 100μM | 100mM | Tocris | 2896/10 |
| Nifedipine<sup>45</sup> | 10, 50, 100μM | 100mM | Tocris | 1075/100 |

**Table 5:** Taqman probes (Thermo Fisher) used for qRT-PCR

| Probe | Cat no. | Exon Boundary |
| --- | --- | --- |
| CD3ε | Mm00599684_g1 | 6-7 |
| Tbp | Mm00446973_m1 | 4-5 |
| Tbp | Mm00446971_m1 | 2-3 |
| Ihh | Mm01259021_m1 | 1-2 |
| Smo | Mm01162710_m1 | 8-9 |
| Gli1 | Mm00494654_m1 | 11-12 |
| Gli1 | Mm00494645_m1 | 2-3 |
| Ulk3 | Mm01151463_m1 | 4-5 |
| CACNA1F | Hs00913730_m1 | 46-47 |
| CACNA1D | Hs00167753_m1 | 46-47 |
| CACNA1S | Hs00163885_m1 | 2-3 |
| CACNA1C | Hs00167681_m1 | 3-4 |
| TBP | Hs00427620_m1 | 2-3 |
| GLI1 | Hs01110766_m1 | 8-9 |

**Table 6:**
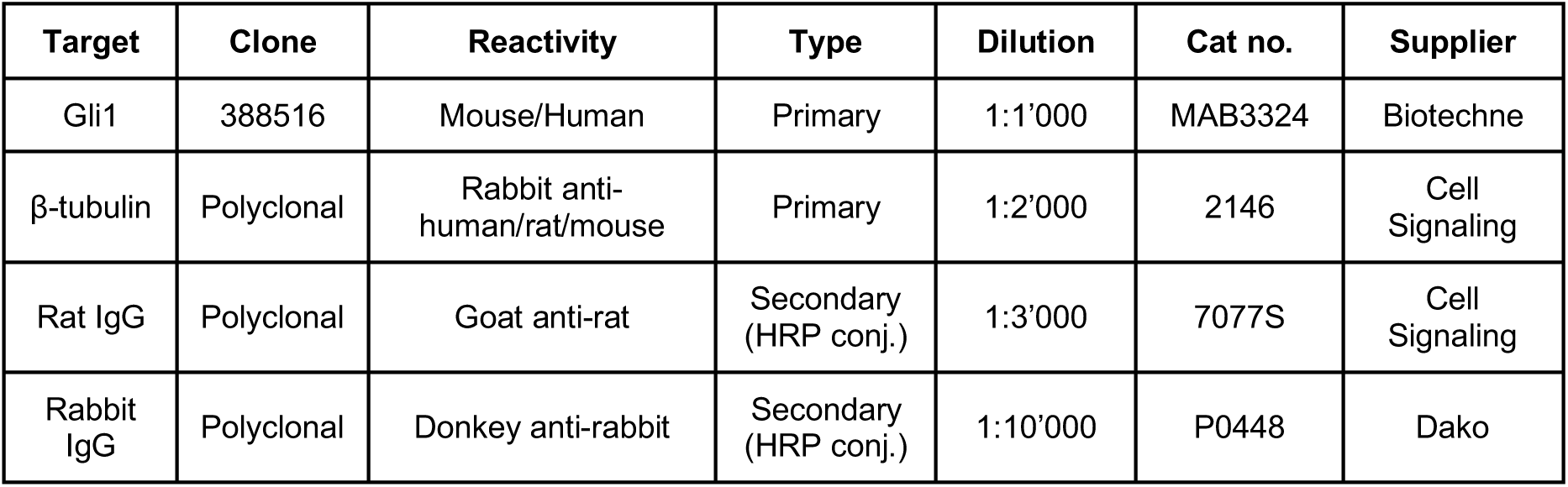
Antibodies used for Western Blotting

### qRT-PCR

Cells harvested for RNA extraction were washed twice in ice-cold PBS, snap-frozen as dry pellets, and stored at -80°C. RNA was extracted using the RNAqeous™-Micro Total RNA Isolation Kit (murine cells) or the Single Cell RNA Purification Kit (Norgen Biotek Corp., cat no. 51800) (human cells) according to the manufacturer’s instructions. RNA concentration was measured with a Nanodrop spectrophotometer (Labtech ND-1000) and samples were stored at -80°C if not used immediately.

Reactions for qRT-PCR were set up using the One-Step qRT-PCR Kit (Thermo Fisher SuperScript III Platinum, cat no. 11732088) according to the manufacturer’s instructions.

Each sample was run in triplicate with *Tbp* and/or *CD3ε* used as housekeeping genes. In addition, each experiment included a non-template control and primer/probes were validated by no RT controls. Samples were run on a QuantStudio 6 Flex Real-Time PCR System (Thermo Fisher). Levels of Gli1 mRNA were determined using probe (Mm00494654_m1) and confirmed in some of the experiments with a second probe (Mm00494645_m1).

Expression of the gene transcript of interest was calculated with the ΔCt method ^46^. The cycle threshold (Ct) value from the gene of interest was subtracted from the housekeeping gene and transformed with a factor of 2^(-ΔCt) to give the fold expression relative to the housekeeping gene.

### Immunofluorescence

Primary human CTLs at day 11 post-stimulation were plated onto glass slides at 4x10^6^/ml and allowed to adhere to the glass for 10min at 37°C in an incubator. Cells were fixed with 4% PFA (16% PFA solution, CN Technical Services, cat no. 15710-s, 1x PBS) for 10min at room temperature. Slides were washed 5 times with PBS and blocked with blocking buffer (PBS + 1% bovine serum albumin (BSA, Sigma, cat no. A3912; 50 g lyophilized powder) + 0.1% TritonX-100 (Alfa Aesar)) for 30min at room temperature. Blocking buffer was aspirated and blocking buffer containing primary rabbit anti-Cav1.4 (1:50, Stratech, Cat no. LS-C94032-LSP) antibody was added. After staining for one hour at room temperature slides were washed 5 times with PBS + 0.1% TritonX-100. Anti-rabbit AlexaFlour 488 secondary antibody (1:400 dilution, Thermo, Cat No. A11034) as well as phalloidin (1:100, Thermofisher, Cat no. A12380) were added in blocking buffer (PBS + 1% BSA + 0.1% TritonX-100) and slides were incubated for 30min light-protected at room temperature. After incubation, slides were washed 5 times with PBS + 0.1% TritonX-100 and stained with Hoechst (Hoechst 33342, trihydrochloride, trihydrate Invitrogen/Fisher, cat no. H3570, 1:30,000 dilution) prepared in PBS for 5min light-protected at room temperature. Slides were washed five times with PBS and mounted with ProLong Diamond Antifade Mountant (Fisher, cat no. P36961).

For conjugate staining, primary human CTLs at day 14/15 post-stimulation (on d0 and d10) were conjugated for 30min at 37°C with P815 target cells at a 1:1 ratio. P815s were pre-stained with 25µM CellTracker Red CMTPX Dye (Thermofisher, Cat no. C34552) for 45min at 37°C and simultaneously pulsed with 1µg/ml CD3 (UCHT1). The staining was comprized of rabbit anti-Cav1.4, phalloidin (1:100, Thermofisher, Cat no. A22287), donkey anti-rabbit 488 and Hoechst 33342, as above.

Confocal spinning disc microscopy was performed on an Andor Dragonfly 500 (Oxford Instruments) and images were processed using Imaris software (Bitplane/Oxford Instruments).

### Western Blot

Cells were lysed in ice-cold RIPA buffer (150mM NaCl, 50mM Tris-HCl pH 8.0, 1mM MgCl2, 2% Triton) with protease inhibitor (Pierce, Cat. No. 88666). Samples were heated at 65°C for 15min and loaded together with a protein standard (Bio Rad, Cat no. 161-0394), onto a NuPAGE 4-12% gradient Bis/Tris Acrylamide gel (Thermo Fisher, Cat no NP0335BOX). PAGE was run in Nu-PAGE MOPS running buffer (Thermo Fisher, Cat no. NP0001). Western Blotting was performed using wet transfer in Nu-PAGE Transfer Buffer (Thermo Fisher, Cat no. NP0006-01) + 10% Methanol (Honeywell, Cat no. 32213-2.5L). Membranes were blocked with 5% (w/v) nonfat dry milk (Marvel Original, Dried Skimmed Milk) in TBS before incubation with primary and secondary antibodies. Membranes were developed with SuperSignal West Pico Plus Chemiluminescent Substrate (Thermo Fisher, Cat no. 34580).

### Retroviral transduction

#### Generation of retrovirus

One day prior to the generation of retrovirus, HEK 293T cells were seeded in a six well plate resulting in 75% confluency the next day. Media was replaced with 2ml fresh DMEM per well prior to transfection with retroviral plasmids. HEK 293T cells were transfected with 1.5µg of packaging plasmid pCL-Eco (generous gift from Gillian Griffiths, Cambridge UK) and 1.9µg pMig vector – either 5’LTR-eGFP-IRES-rThy1.1(CD90.1)-3’LTR (referred to as “empty vector”), 5’LTR-Smo-C terminal eGFP fusion-STOP-IRES-rThy1.1(CD90.1)-3’LTR (referred to as “SmoM2”) or 5’LTR-Ihh-STOP-IRES-rThy1.1(CD90.1)-3’LTR (referred to as “Ihh”) – prepared in Opti-MEM™ media (Gibco, Cat no. 31985070). Concurrently, a 1:25 dilution of Lipofectamine™ 2000 (Invitrogen, Cat no. 11668019) was prepared in Opti-MEM™ media. The DNA-Opti-MEM™ solution was mixed with the Lipofectamine™2000-Opti-MEM™ solution at a ratio of 1:1 and incubated for 5min at room temperature after which 400µl of the mixture was added drop-wise to the HEK-293T cells. Media was replaced 18h after transfection and collected 48h post transfection. The retroviral supernatant was passed through a 0.45µm PVDF membrane filter (Merck Millipore, Cat no. SLHVM33RS) for immediate use.

#### Retroviral transduction of CD8^+^ T cells

Naïve CD8^+^ T cells were isolated from C57BL/6 spleens and stimulated with plate bound anti-CD3*ε*/CD28 antibodies as described previously. After 24h of stimulation, T cell media was partially withdrawn and replaced by retroviral supernatant in a 1:2 ratio supplemented with protamine sulphate (Sigma, Cat no. 1101230005) at a final concentration of 9µg/ml and IL-2 to maintain the standard CD8^+^ T cell polarization conditions as described above. Cells were then centrifuged at 1800rpm for 15min at 32°C and subsequently placed in a humidified cell culture incubator. 48h cells were centrifuged to wash off retroviral supernatant and cultured after that in complete RPMI + IL-2.

#### Magnetic sorting of retrovirally transduced cells

For downstream applications such as qRT-PCR, retrovirally transduced cell populations were enriched for transduced cells measured by the presence Thy1.1 expression. T cell populations were subjected to magnetic sorting at day 3 or 4 after initial stimulation. Thy1.1^+^ cells were positively enriched using the EasySep™ PE Positive Selection Kit (Stem Cell Technologies, Cat no. 17684) or EasySep™ Release Mouse PE Positive Selection Kit (Stem Cell Technologies, Cat no. 17656) according to the manufacturer’s instructions. Cells were labelled with a PE-conjugated Thy1.1 antibody (Clone HIS51, BD, Cat No. 12-0900-81) at a final concentration of 0.25mg/ml in MACS buffer. All steps were performed in MACS buffer and all incubations were performed at room temperature. PE positive cells were enriched using an EasyEights™ magnet (Stem Cell Technologies, Cat no. 18103). Three rounds of separation were performed in total and cells were washed in complete media prior to maintenance in culture with complete RPMI + IL-2. Purity of the sort was routinely over 95% as measured by FACS.

### Nucleofection of CTLs

The GFP-VIVIT plasmid was a gift from Anjana Rao (Addgene Plasmid # 11106; http://n2t.net/addgene:11106; RRID:Addgene_1106) which contains a fusion of MAGPHPVIVITGPHEE to the N-terminus EGFP, resulting in selective inhibition of the interaction between calcineurin and NFAT^16^. The IκBα dominant negative (“IκB DN”) plasmid was a generous gift from Felix Randow (Laboratory of Molecular Biology, Cambridge). The plasmid contains an IκBα which is resistant to proteosomal degradation due to two amino acid substitutions (S32→A32 and S36→A36) resulting in a dominant-negative phenotyping inhibiting NFkB induced transcription^17^.

Splenocytes from one OT-I Rag2^-/-^ mouse were stimulated with 10nM Ova (SIINFEKL) peptide for 48h and subsequently washed daily (resuspended at 1x10^6^ cells/ml). On day 6 CTLs were subjected to lympholyte treatment (Cedarlane, CL5035) per the manufacturer’s instructions. On day 7 cells were nucleofected using the Amaxa Mouse T Cell Nucleofector Kit (Lonza, Cat no. VVPA-1006) per the manufacturer’s instructions with 5x10^6^ cells and 1.5µg plasmid DNA per cuvette. Cells were electroporated using a Nucleofector 2b Device (Lonza) on programme X-001. Cells were restimulated 18h post nucleofection on day 8.

### ImageStream analysis of NFkB nuclear translocation

On day 8, 18h after nucleofection, CTLs were counted and plated onto a 24-well plate pre-coated overnight with 2.5µg/ml anti-CD3*ε* antibody. After 1h of stimulation, cells were harvested and washed once in PBS prior to staining with eFluor780 for 10min light-protected at room temperature. Cells were subsequently washed in perm wash buffer – made up of 2% FCS, 0.1% Sodium Azide, 0.1% Triton X-100 (Alfa Aesar, Cat no. A16046.AE) in PBS prior to fixation in 4% PFA for 10min light-protected at room temperature. Cells were washed twice in perm wash buffer before staining in 100µl perm wash buffer containing anti-p65 APC antibody (Table 4) for 30min light-protected at room temperature. Cells were washed twice in perm wash buffer, subsequently resuspended in 50µl perm wash buffer and transferred to a 1.5ml Eppendorf tube. DAPI was added (0.5mg/ml) immediately prior to acquisition on an Amnis ImageStream (MilliporeSigma) imaging cytometer. Data was analysed with IDEAS Software (MilliporeSigma).

### Analysis of Chip-Seq data

Coverage bigwig files were downloaded from GEO (GSE54191). The Integrative Genomics Viewer (IGV) was used for coverage analysis^47^.

## DATA AVAILABILITY

This study includes no data deposited in external repositories.

## ACKNOWLEDGEMENTS

We thank all members of the de la Roche lab for constructive comments on the manuscript. Special thanks and gratitude go to Ellie Pryor, Nicky Jacobs and Gemma Cronshaw from the BRU for expert animal care; Fadwa Joud from the Microscopy core for assistance in confocal microscopy and the flow cytometry core for assistance with cell sorting. We thank Rose Zamoyska for advice on CRISPR experiments in primary T cells, Fanni Gergely for help with reagents, and Gillian Griffiths for advice on the manuscript.

## FUNDING

This work was supported by Cancer Research UK (MdlR (A22257): CK, VC, FB, CC); Sir Henry Dale Fellowship jointly funded by the Wellcome Trust and the Royal Society (MdlR (WT107609): LMOB); JH is undertaking a PhD funded by the Cambridge School of Clinical Medicine, Frank Edward Elmore Fund and the Medical Research Council’s Doctoral Training Partnership (award reference: 1954837).

## AUTHOR CONTRIBUTION

MdlR and JH conceived the project and designed the experiments. JH executed most experiments. CK performed experiments with human CD8^+^ T cells with assistance from FB. LMOB and VC performed experiments and helped with technical expertise, and Chandra Chilamakuri analysed Chip-Seq data. JH and MdlR analyzed the results and wrote the manuscript.

## CONFLICTS OF INTEREST

The authors declare that they have no conflict of interest.

## Notes

### Competing Interest Statement

The authors have declared no competing interest.

