## Supplemental Figures for "Non-canonical Hedgehog signaling through L-type voltage gated Ca^2+^ channels controls CD8^+^ T cell killing"

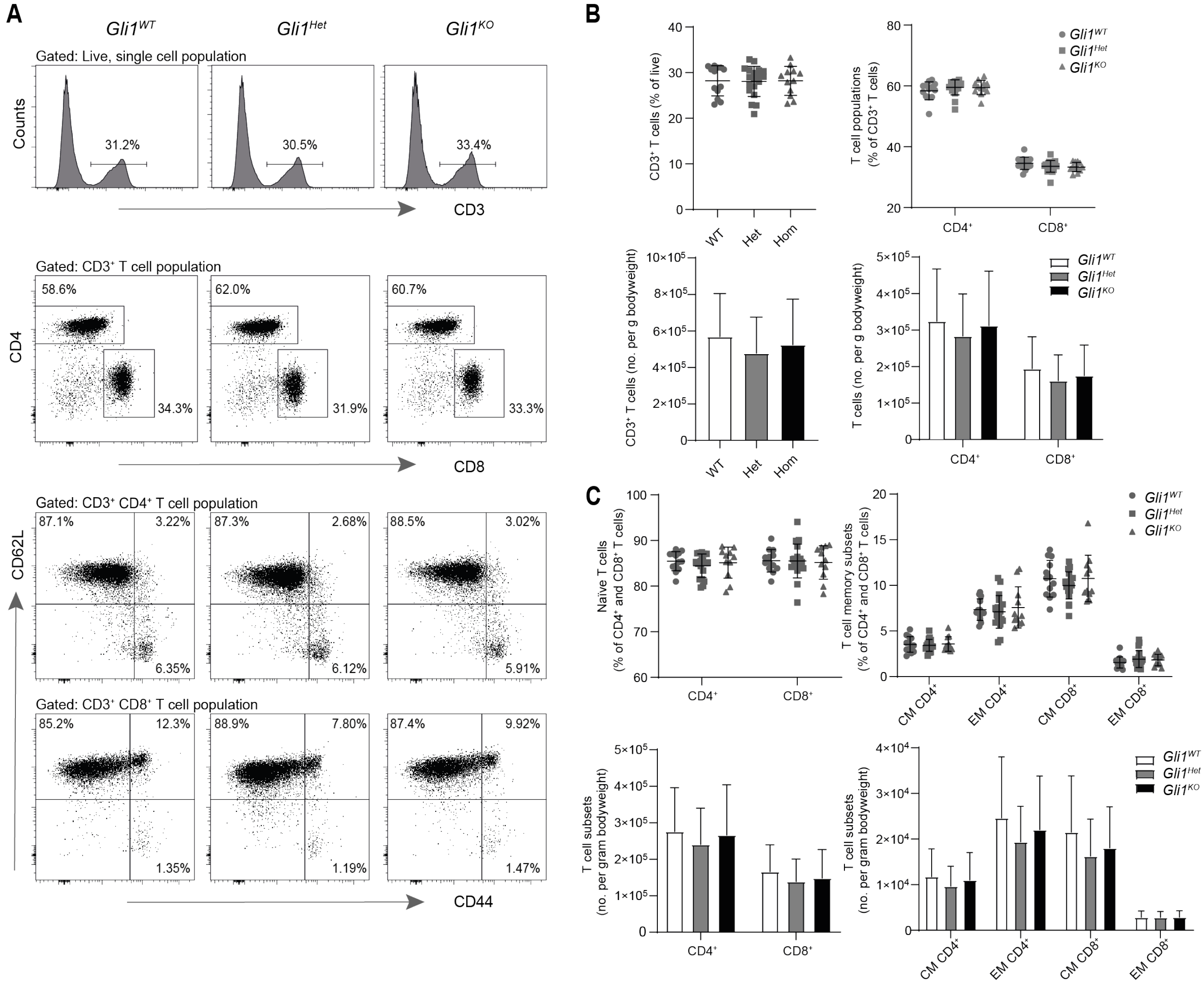


**Suppl. Figure 1: *Gli1$^{KO}$* mice have phenotypically normal peripheral T cells.** Splenocytes were isolated from *Gli1$^{WT}$*, *Gli1$^{Het}$*, and *Gli1$^{KO}$* mice. (**A**) Splenocytes were stained for FACS analysis of T cells ($CD3^+$, top panel), $CD4^+$ and $CD8^+$ T cells (middle panel), and naïve ($CD62L^+$, $CD44^-$), central memory ($CD62L^+$, $CD44^+$), and effector memory ($CD62L^-$, $CD44^-$) subsets (bottom two panels). Representative FACS plots for WT, Het, and KO mice are shown. (**B and C**) Quantitative analysis of percentages and cell numbers from stainings in (A). Every circle (WT), square (Het), and triangle (KO) represents a single mouse. Error bars indicate SD.

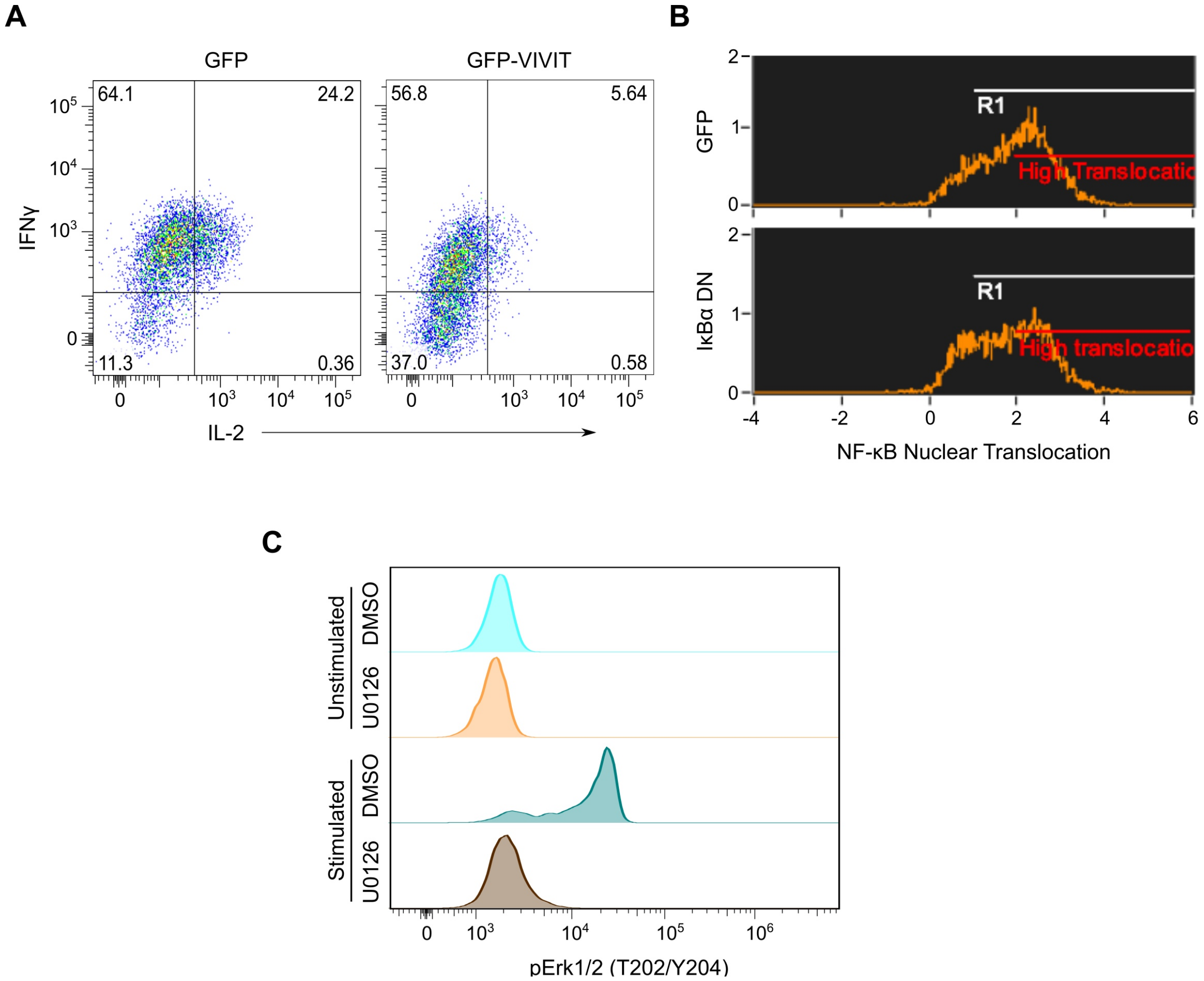


**Suppl. Figure 2: Validation of TCR signaling inhibitors.**
(**A**) CTLs were nucleofected with GFP or GFP-VIVIT on day 6. Cells were restimulated with PMA/Ionomycin on day 7 prior to intracellular flow cytometric staining for IL-2 and IFNγ. FACS plots are gated on $GFP^+$ cells. (**B**) CTLs were nucleofected with GFP or Dominant Negative IκBα (DN IκBα) on day 6. Cells were restimulated with anti-CD3 on day 7 for 1h prior to ImageStream analysis. (**C**) Naïve $CD8^+$ T cells were isolated from spleens and peripheral lymph nodes of $Rag2^{-/-}$ OT-I mice and stimulated with cross-linked soluble anti-CD3/CD28 in the presence of 10μM U0126 or carrier control prior to analysis by flow cytometry of phosphorylated Erk1/2 (pErk1/2). (**A-C**) One representative experiment of two to three independent experiments is shown.

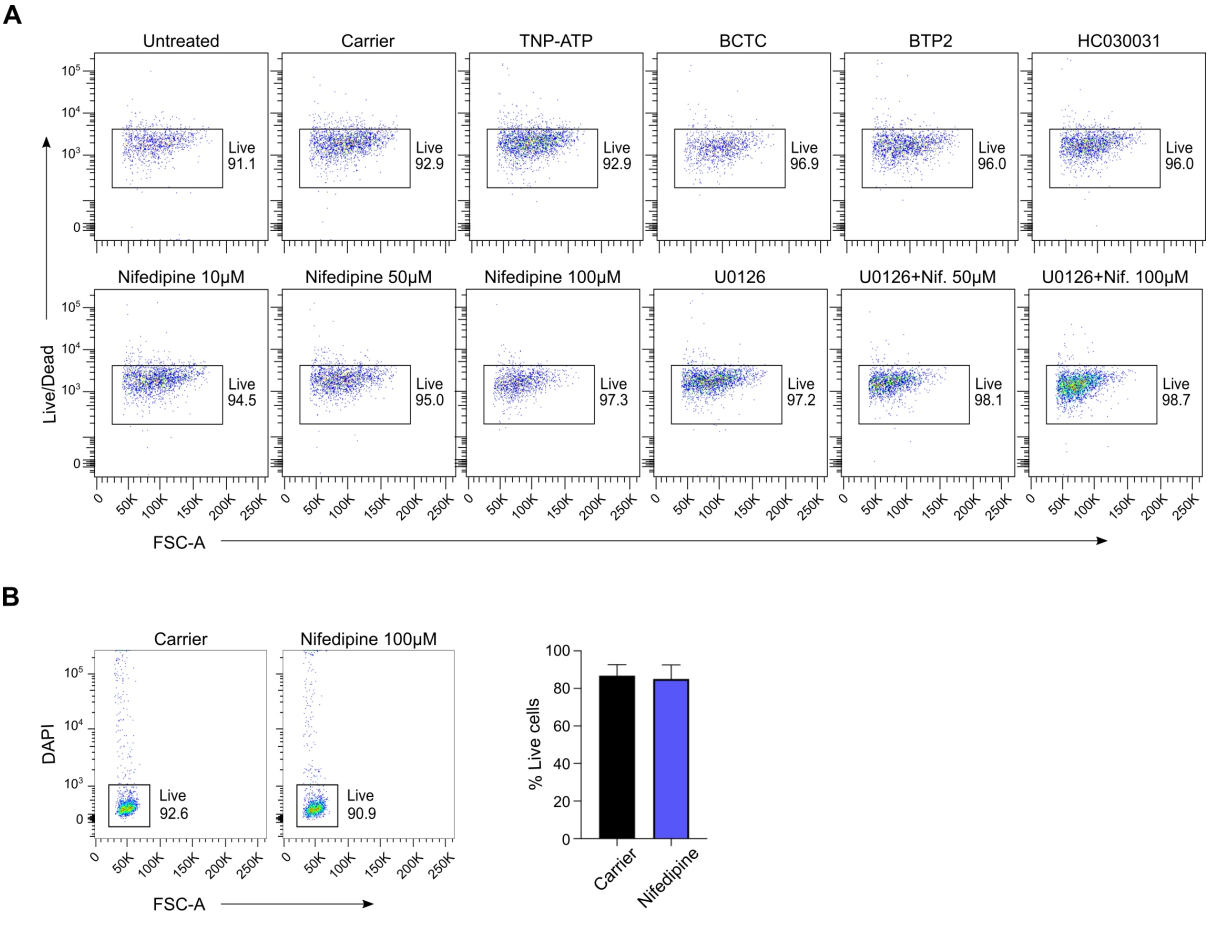


**Suppl. Figure 3: Cell viability of mouse and human CD8$^+$ T cells upon Ca$^{2+}$ channel blockade**

(**A**) CTLs were restimulated with plate-bound anti-CD3 in the presence of the indicated inhibitors or carrier control for 15h before flow cytometric analysis for cell viability. (**B**) Naïve human CD8$^+$ T cells were isolated from PBMCs of Healthy Donors and stimulated with human T-activator CD3/CD28 beads for 3 days before flow cytometric analysis for cell viability.

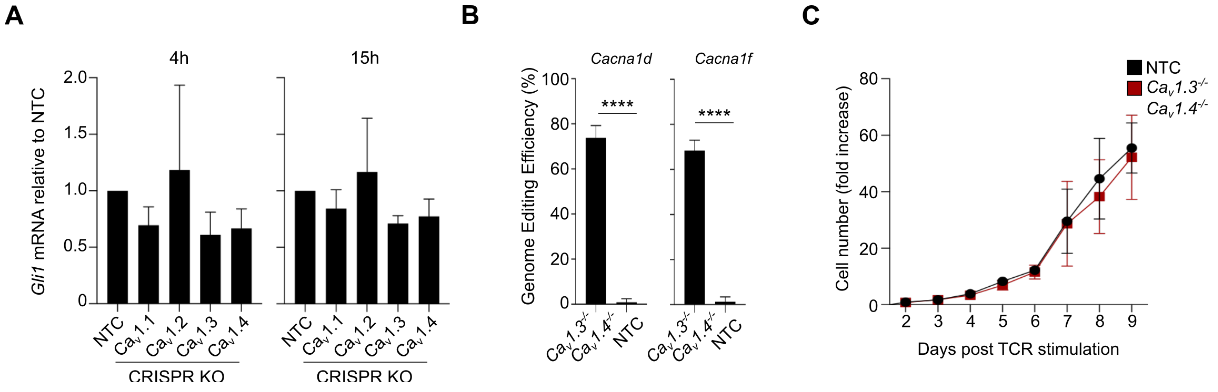


**Suppl. Figure 4: Characterization of $Ca_v1$ CRISPR knockouts**

(**A**) CTLs were electroporated with RNP complexes at day 2 post stimulation to generate *$Ca_v1.1^{-/-}$, $Ca_v1.2^{-/-}$, $Ca_v1.3^{-/-}$, $Ca_v1.4^{-/-}$* (KO) or non-targeting control (NTC) CTLs. CTLs were restimulated on day 8/9 with plate-bound anti-CD3ε for qRT-PCR analysis. n=3 independent experiments. Data is normalized to *CD3ε* as a reference gene. Similar results were obtained when *Tbp* was used as a reference gene. (**B**) *$Ca_v1.3^{-/-}$* and *$Ca_v1.4^{-/-}$* CRISPR double KO CTLs were generated as described previously. PCR amplification of the edited loci was performed prior to a T7EI mismatch cleavage assay to quantify genome editing efficiency. n=3 independent experiments. Error bars indicate SD. p values were calculated using an unpaired two-tailed Student's t test. **** indicates $p<0.0001$. (**C**) *$Ca_v1.3^{-/-}$* and *$Ca_v1.4^{-/-}$* CRISPR double KO CTLs and NTC CTLs were generated as described previously. Fold expansion in cell numbers after CRISPR performed at day 2 post-stimulation is shown. n=3 independent experiments.

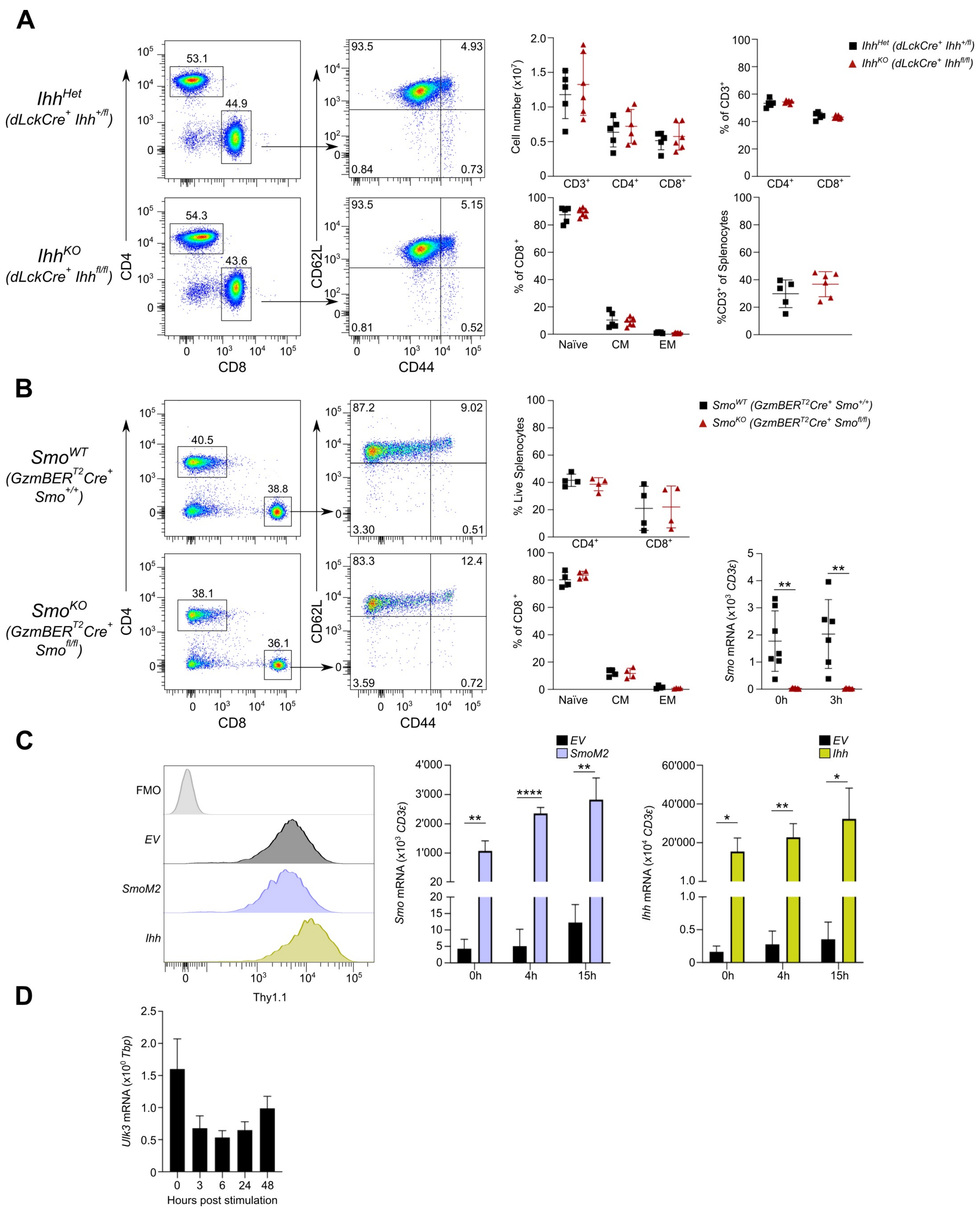


**Suppl. Figure 5**

**Suppl. Figure 5: Characterization of *dLckCre⁺ Ihh^KO*, *GzmBER^T2Cre⁺ Smo^KO* mice and retroviral transduction constructs**

(**A**) Splenocytes were isolated from *dLckCre*$^{+}$*Ihh*$^{+/fl}$ (*Ihh*$^{Het}$) and *dLckCre*$^{+}$*Ihh*$^{fl/fl}$ (*Ihh*$^{KO}$) mice and subjected to flow cytometric phenotypic analysis. Representative FACS plots are shown on the left. Quantification of relative percentages, cell numbers and steady-state memory phenotype (naïve (CD62L$^{+}$, CD44$^{-}$), central memory (CM, CD62L$^{+}$, CD44$^{+}$), and effector memory (EM, CD62L$^{-}$, CD44$^{-}$) shown on the right. Every square represents one HET, and every triangle one individual KO mouse. (**B**) Splenocytes were isolated from *GzmBER*$^{T2}$*Cre*$^{+}$*Smo*$^{+/+}$ (*Smo*$^{WT}$) and *GzmBER*$^{T2}$*Cre*$^{+}$ *Smo*$^{fl/fl}$ (*Smo*$^{KO}$) mice and subjected to flow cytometric phenotypic analysis. Quantification of relative percentages and steady-state memory phenotype shown on the right. Every square represents one WT, and every triangle one individual KO mouse. CTLs were generated from these mice and restimulated at day 8/9 for qRT-PCR analysis of Smo. (**C**) CD8$^{+}$ T cells were retrovirally transduced with constructs encoding empty vector (EV), SmoM2 or Ihh, respectively. Representative flow cytometry plots of sorted, transduced (Thy1.1$^{+}$) cell populations shown (left panel). Sorted, transduced CTLs were restimulated at day 8/9 for qRT-PCR analysis of Smo (middle panel) or Ihh (right panel), respectively. n=3 independent experiments. (**D**) Naïve CD8$^{+}$ T cells were isolated from C57BL/6 mice and stimulated with plate-bound anti-CD3/CD28 antibodies for the indicated times before qRT-PCR analysis for *Ulk3* mRNA. Data is normalized to *Tbp* as a reference gene. (**B/C**) Data is normalized to *CD3*$\varepsilon$ as a reference gene. Similar results were obtained when *Tbp* was used as a reference gene. Error bars indicate SD. p values were calculated using an unpaired two-tailed Student’s t test. * indicates $p<0.05$, ** indicates $p<0.01$, **** indicates $p<0.0001$.
